# Early vertebrates were not slow: rapid life histories in Devonian agnathans

**DOI:** 10.64898/2026.04.26.720944

**Authors:** Nanako Okabe, Daniel Pauly

## Abstract

Life history strategies such as rapid growth and high population turnover rates observed in vertebrates have been thought to have emerged relatively late in evolution. However, very little direct evidence exists at the species level for early vertebrates. In this study, a large fossil collection of over 450 specimens of *Protaspis* spp, heterostracan agnathans from the Cottonwood Canyon Formation at Beartooth Butte, Wyoming, from the Early Devonian, was analyzed. Morphological observations, analysis of bone plate completeness, and length-frequency analyses using ELEFAN—commonly used in recent fish studies—were applied to reconstruct growth rates, cohort structure, and ontogenetic processes. *Protaspis* specimens exhibited a continuous growth series from juvenile to adult stages, and a clear cohort structure was identified from the length-frequency distributions. The ELEFAN analysis suggested a life-history characterized by rapid growth and a short life span, and these features remained consistent in subset analyses restricted by species or locality, confirming the robustness of the estimates. Furthermore, the integration of the dermal bone plates progressed during the late stages of ontogeny, revealing that the rapid growth during the juvenile stage preceded the completion of this defensive structure. Comparisons of their growth parameters with those of extant fishes show that *Protaspis* does not align with slow-growing, long-lived “living fossil” taxa, but instead clusters with small-bodied, fast-growing species. These findings suggest that life-history strategies involving rapid growth and high population turnover were already established in early jawless vertebrates, much earlier than previously assumed.

## Introduction

Life history strategies, including growth rate, timing of maturation, and population turnover, form a fundamental link connecting ontogeny, population dynamics, and ecosystem structure in vertebrates (Arendt, 1997; de Roos, 2020; Stearns, 1977, 1998). In extant fish, life histories span a broad continuum, ranging from rapidly growing, early-maturing, high-mortality species (e.g., the gobies, Familiy Gobiidae) to taxa exhibiting slow growth, long lifespans, and delayed reproduction (e.g., the sturgeons, Family Acipenseridae). In contrast, because direct evidence from fossil populations is scarce, the ecology of early vertebrates is often inferred through comparisons with genetically closely related living “living fossils” and heavily armored Paleozoic fish, including early gnathostomes and placoderms. Thus, it is generally assumed that they adopted a uniformly conservative strategy characterized by slow growth and low population turnover. As a result, despite maintaining ecological dominance in marine ecosystems for tens of millions of years before the emergence of jawed vertebrates, the ecological diversity and population dynamics of early vertebrates remain poorly understood.

The primary barrier to reconstructing the life history of early vertebrates is the lack of developmental data in the fossil record. The skeletal tissues of immature individuals are rarely preserved, and particularly in Paleozoic fish, the early stages of ossification are rarely fossilized (Chevrinais et al., 2015). Consequently, developmental stages are often inferred from a very small number of juvenile specimens, introducing significant uncertainty into growth trajectories and life history interpretations (Cloutier, 2010). While fossil ontogeny holds the potential to reveal developmental patterns and evolutionary processes impossible to elucidate in extant taxa, such datasets have been extremely limited. Only very recently, a large-scale ontogenetic study based on over 3,500 specimens of the Devonian agnathan *Euphanerops longaevus* demonstrated that early vertebrate development was more highly organized than previously anticipated. Furthermore, it proved the utility of population-level fossil data, such as inferring ecology from populations (Chevrinais et al., 2023). In contrast, equivalent studies on Heterostraci remain limited to a small sample of fewer than 10 specimens, primarily from the Family Cyathaspididae (Greeniaus & Wilson, 2003) and are confined to descriptions of basic body ossification processes.

Here, we focus on the heterostraci agnathan genus *Protaspis* from the Early Devonian Beartooth Butte formation in Wyoming (USA). It is represented by over 600 exceptionally well-preserved specimens housed in the Field Museum of Natural History and the University of Kansas Natural History Museum. These specimens originate from a single depositional environment, interpreted as sediments of a warm brackish water channel located near the terminus of a river system (Fiorillo, 2000; Lamsdell & Legg, 2010; Lamsdell & Selden, 2013). This represents an assemblage of mass mortality, likely capturing a nearly simultaneous snapshot of the living population. This assemblage uniquely preserves a continuous growth series from small juveniles to adults, as it is covered by multiple hard dermal plates, an extremely rare occurrence in agnathans (Denison, 1967). While this material includes specimens historically assigned to several species, they are treated as a single population due to their highly similar growth patterns and sizes, and for the purposes of growth and population dynamics analysis. Nevertheless, the scale and completeness of this collection enabled the application of Electronic Length-Frequency Analysis or ‘ELEFAN’ (Pauly & David, 1981), a quantitative, but non-parametric and robust method developed in fisheries science to estimates the parameters of asymptotic growth curves from length-frequency data of fish and marine invertebrates.

The growth parameters obtained here allow comparison with extant fishes. Furthermore, combining the growth curves with detailed observations on dermal plate development provides a robust framework for reevaluating the growth process from larvae to adults, life history strategies, and ecological roles in early jawless vertebrates.

## Results

### Morphological description

The morphology of *Protaspis* spp. changes with growth, and its individuals also show several stages. First, based on basic observations of the specimens: Small *Protaspis*, considered to be in the juvenile stage with a dorsal plate around 10 mm, only show a dorsal plate and a tail-like structure (Figure 1A, B, C). Subsequently, when the dorsal plate reaches 15 mm, the rostrum, orbital plate, branchial plate, and a scaled tail become identifiable (Figure 1 D, E, F). Its morphology is rounded when small, becoming sharper overall with increasing size. By the time the dorsal plate reaches approximately 50 mm, all plates are fully formed and closely packed together (Figure 1 G, H, I). Furthermore, while *Protaspis* is composed of multiple plates, many small and ephemeral, the dorsal or ventral plates cover most of the body and are stably preserved. Therefore, their length and frequency were examined (Figure S1). The results showed two distinct peaks in a histogram of dorsal plate lengths: one at approximately 15 mm and another at approximately 90 mm. The bimodal size distribution is consistent with the occurrence of at least two distinct cohorts within a single population. Considering this is a snapshot fossil assemblage, this likely indicates *Protaspis* juvenile and adult stages inhabited the same area, and that reproduction and recruitment occurred once a year within a well-defined season.

**Figure 1.**
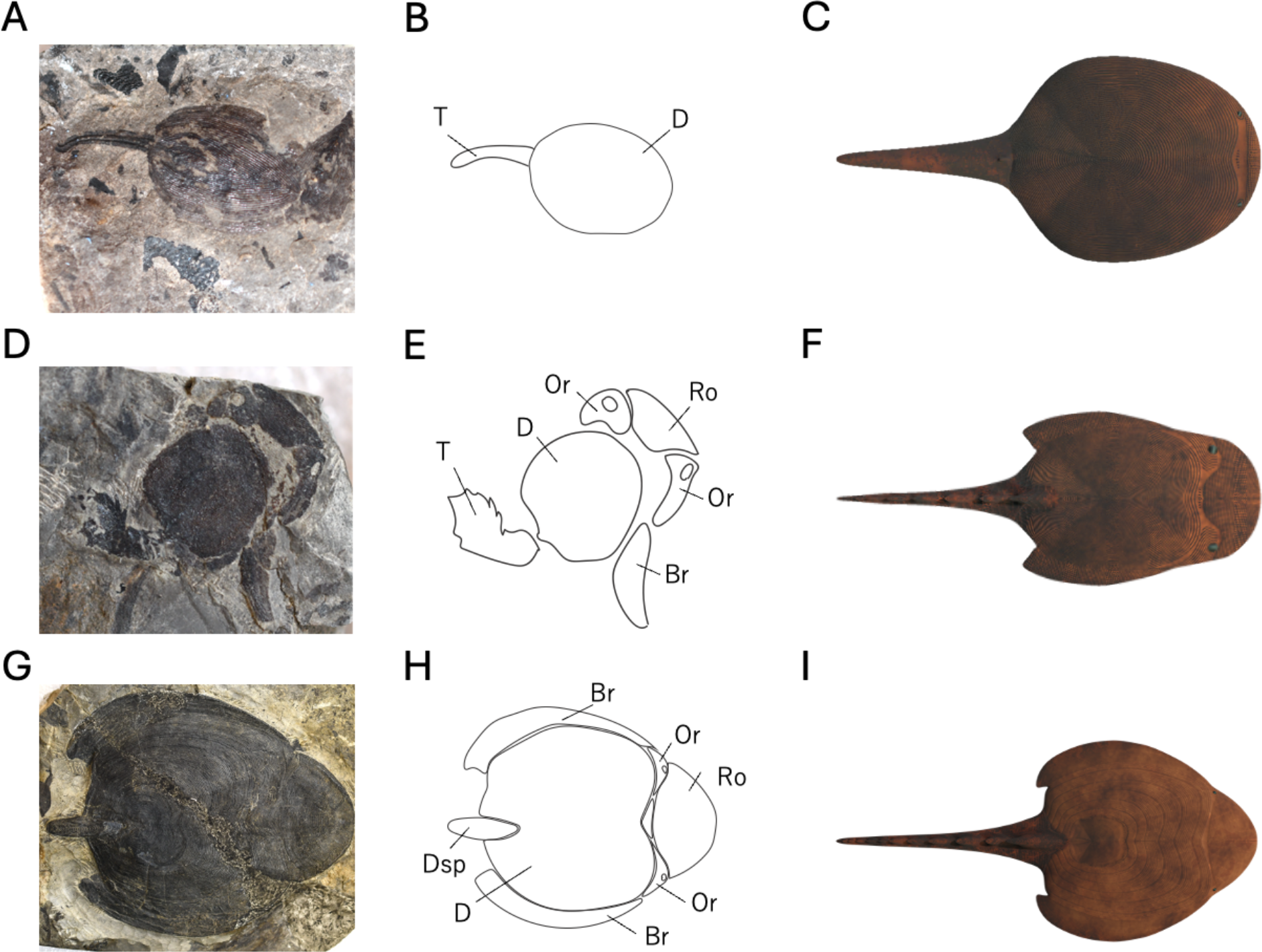
Fossil specimens and reconstructions of *Protaspis* at each growth stage. (A) Fossil specimen at the smallest stage. PF5159, Dorsal plate length=10.673mm. (B) Diagram of A. (C) Generalized reconstruction model of A. (D) Fossil specimen of the intermediate stage. PF4958, Dorsal plate length=15.134mm. (E) Diagram of D. (F) Generalized reconstruction model of D. (G) Fossil specimen of the enlarged form. PF4334, Dorsal plate length =53.61mm. (H) Diagram of G. (I) Generalized reconstruction model diagram of G. Symbols in the diagrams represent: D = Dorsal plate, T = Tail, Ro = Rostrum, Or = Orbital, Br = Branchial plate, Dsp=Dorsal spine plate.

### Plates completeness analysis

As growth altered the preservation rate of plates and their mutual adhesion, we attempted to clarify growth stages based on their integrity. Results from the main dataset (cw), which includes several species, showed that while the dorsal plate maintained high integrity throughout, the preservation rate of other plates increased around the time the dorsal plate reached 60-80 mm (Figure 2C). This pattern was consistently observed in independent subset (cwtr, ss, sstr) analyses, including datasets restricted to a single species *Cosmaspis (Protaspis) trasnversa* indicating that the result is robust to taxonomic composition (Fig. S6). This likely indicates that as growth progressed, the plates forming the outer layer of *Protaspis* became rigidly fused and less brittle. Growth continues beyond approximately 70 mm, but the articulations become rigid, exhibiting a more stable morphology. This suggests that specimens below around 70 mm can be considered juveniles, prioritizing growth rate over stability, while those above around 70 mm exhibit the stable morphology characteristic of adults.

**Figure 2.**
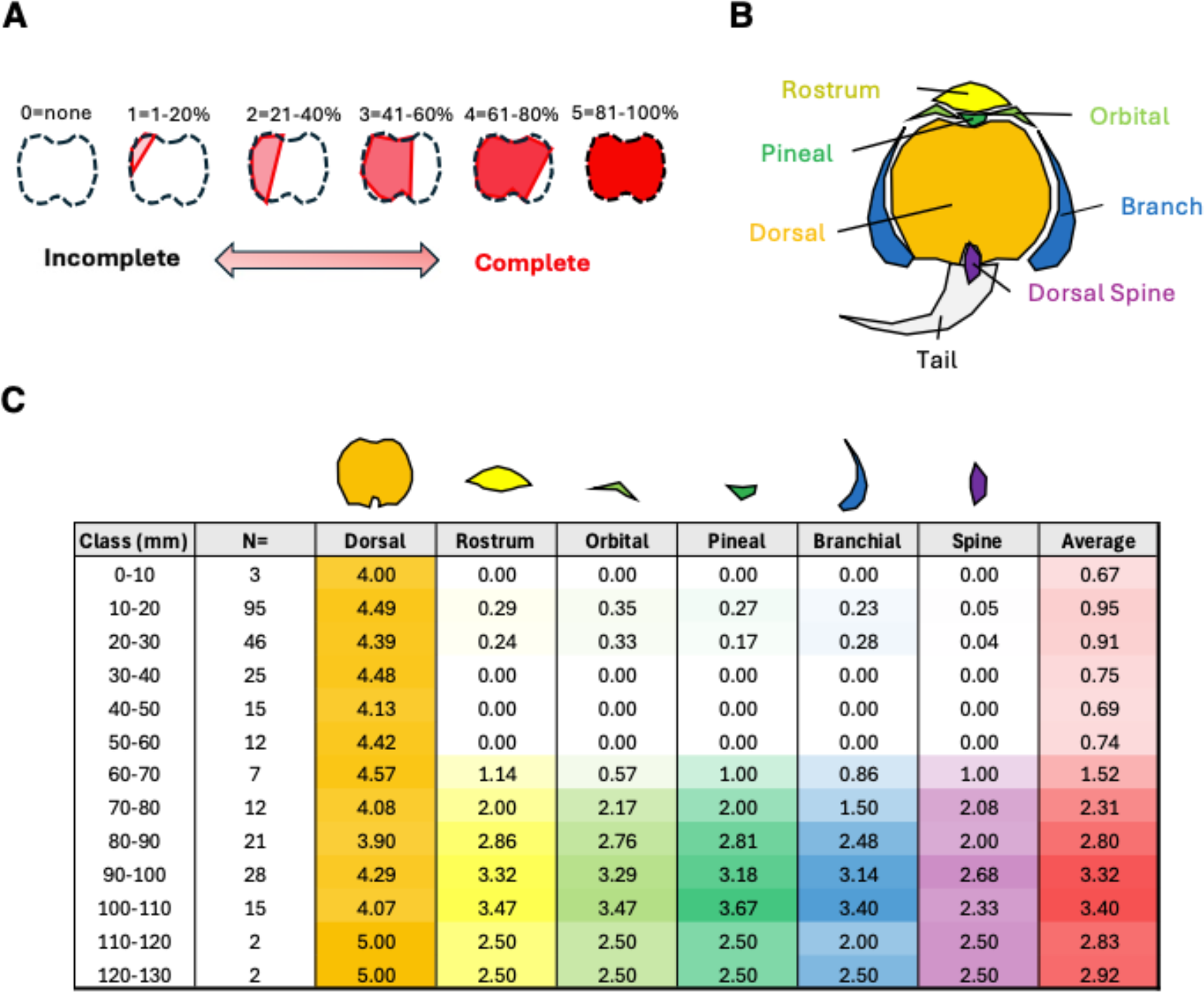
Schematic diagram of Plate Completeness scoring, illustrations of each plate name and Plate completeness score of each plate of *Protaspis*. (A) Score of plate completeness. 0=none, 1=1-20%, 2=21-40%, 3=41-60%, 4=61-80%, 5=81-100%. (B) Plate: Rostrum, Orbital, Pineal, Dorsal, Branch, Dorsal spine. (C) Select specimens possessing a dorsal plate and represent them with a color-coded heatmap based on the average plate completeness score of the other plates (Rostrum, Orbital, Pineal, Branch, Dorsal Spine). The average score is the mean score of the other plates, and n = x represents the number of samples per size bin for each dorsal plate. Size bins are designated as greater than or equal to and less than or equal to. For example, “0–10” denotes 0 mm or greater but less than 10 mm.

### ELEFAN analysis

Analysis of length-frequencies of the dorsal and ventral plates of 455 individuals of the genus *Protaspis* from Cottonwood Canyon revealed a clear cohort structure suitable for ELEFAN analysis, which fits the von Bertalanffy Growth Function (Von Bertalanffy, 1938) to length-frequency (L/F) data. As this analysis requires the available L/F data to be grouped in histograms, the effect of the histograms class interval or ‘bin’ size on growth parameter estimation intervals was assessed by varying the bin size from 5.0 mm to 20.0 mm in 0.1 mm increments and the resulting estimates of K were plotted against interval size. These K values did fluctuate with interval size, however, since it was apparent that the K values remained stable within the range of 8.7 to 11.6 mm (Figure 3A), it was defined as the stable region, and we used an estimated parameter K = 0.95 ± 0.015 (SE) and L_∞_ = 140. Hence the growth of *Protaspis* (main dataset: cw) can be represented by

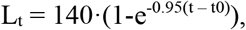

where L_t_ is the length at age t (in years), 140 is the asymptotic length (L_∞_; in mm), 0.95 is K, expressing the rate at which L_∞_ is approached (in year^-1^) and t_0_ (in years) adjust a growth curve to an absolute age scale, and is here assumed to be zero.

**Fig. 3.**
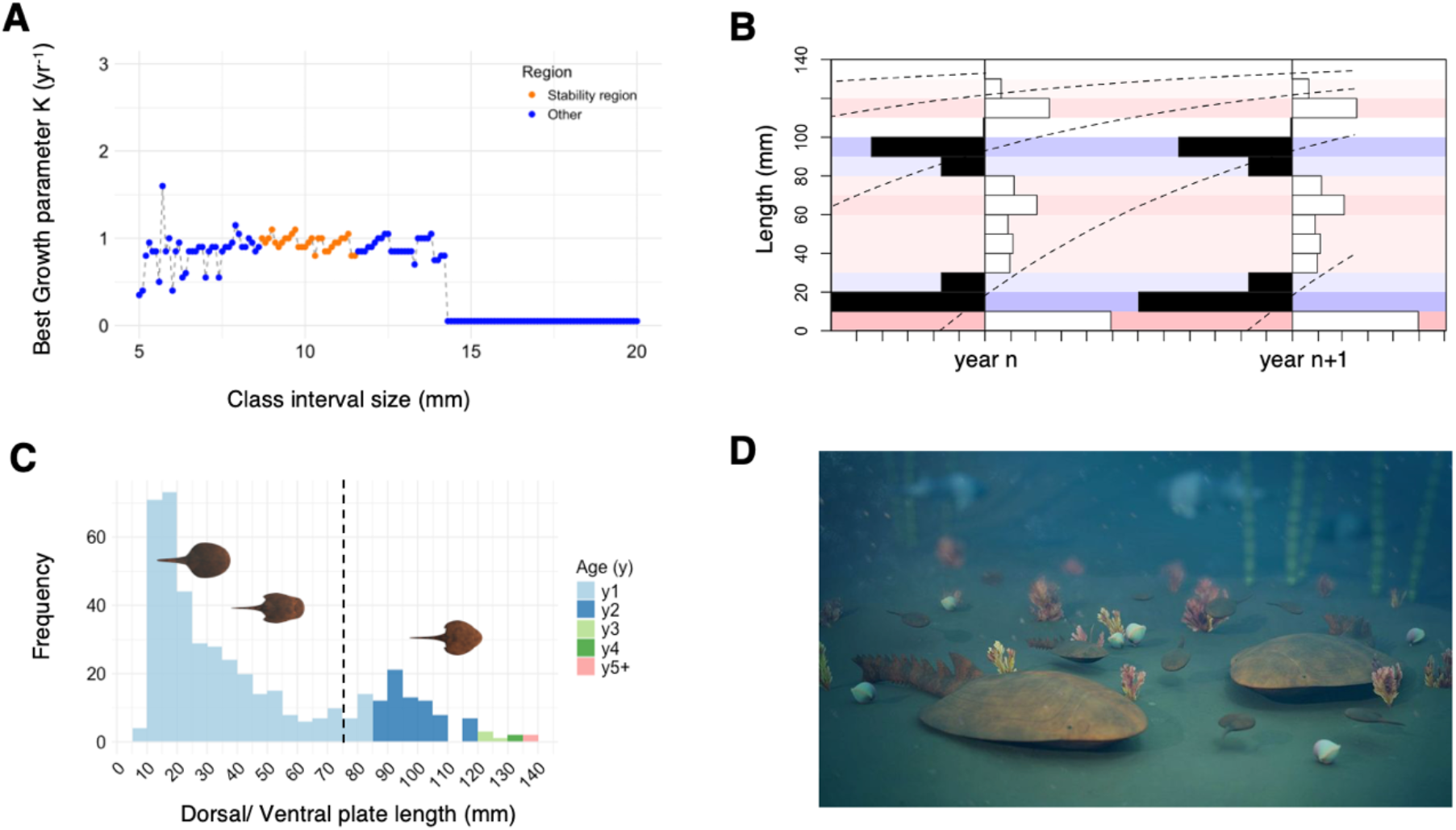
Age Estimation by Size Using ELEFAN Analysis, Associated Parameter Settings and Habitat Restoration Diagram. (A) Search for the optimal K value (BestK) within the Best Growth constant (K) range. The X-axis represents the bin size (mm) of the class interval, and the Y-axis shows the corresponding BestK values. Stability was judged when the interval bin size was 8.7-11.6 mm (Bin size dividing the asymptotic length L_∞_ = 140 mm into 12–16 equal parts). The Mean K in this range was 0.952 year^-1^, with a standard error of 0.015. (B) ELEFAN plot using estimated parameters. *Protaspis* population data was treated as two years of data, and ELEFAN was run using the estimated L_oo_ and K. The dotted line shows the estimated average size per age. (C) Age-specific size data for *Protaspis* population structure. 1y = 0–1, 2y = 1–2, 3y = 2–3, 4y = 3–4 years. X-axis: Dorsal/ventral plate length (mm). Y-axis: Frequency. Draw a gray dotted line at the plate length of 75 mm, which marks the transition point between Juvenile and Adult as determined by plate completeness data, and illustrate the morphological changes before and after this point. (D) A reconstructed habitat map created using data from plate completeness and ELEFAN. Illustrations by Franz Anthony (They inhabit the same area from Juvenile to Adult, with Juveniles being more numerous, and their numbers decreasing as they mature.

This yielded a growth curve that faithfully followed the modal progression of the length frequency data (Figure 3B). The resulting VBGF provided relative ‘ages’ at length estimates and thus could be used to assign individuals to specific cohorts. Length frequency histograms, color-coded by estimated age group, show clear cohort progression consistent with the estimated growth trajectory (Figure 3C).

To assess the robustness of these results, the same analysis was performed on multiple subsets of the dataset, including all specimens (subset: cwtr) and only *C. transversa* (subset: ss) from S side of Cottonwood Canyon locality and a subset restricted to *C. transversa* from Cottonwood Canyon (subset: sstr). The growth parameters obtained from these analyses were generally consistent with the results from the full Cottonwood Canyon dataset. Detailed results of the subset analyses are shown in Supplementary Figure S7. From these population structures, the Juveniles appears to be more numerous, suggesting it shares the same habitat as the Adults (Figure 3D).

### Comparing with extant fishes

Using the VBGF parameters L_∞_ and K estimated for *Protaspis*, comparisons were made with extant species based on ‘auximetric plots,’ i.e., plots of log(K) vs log(L_∞_) (Cury & Pauly, 2000). The parameters for extant species were obtained for the global online encyclopedia of fish called rFishBase (Froese & Pauly, 2026) using re rFishBase routine of Boettiger et al. (2012).

Re-expressing the above asymptotic length of *Protaspis* (140 mm; based on dorsal/ventral plates) as total body length (including the tail) leads to L_∞_ ≈ 28 cm, while K remains = 0.95 year^-1^. The highest density of fish species from rFishBase occurred at L_∞_ =40.5 cm and K = 0.25 year^-1^ (Figure 4A). This suggests the *Protaspis* grew faster and belongs to a group that did not reach large body sizes.

**Fig. 4.**
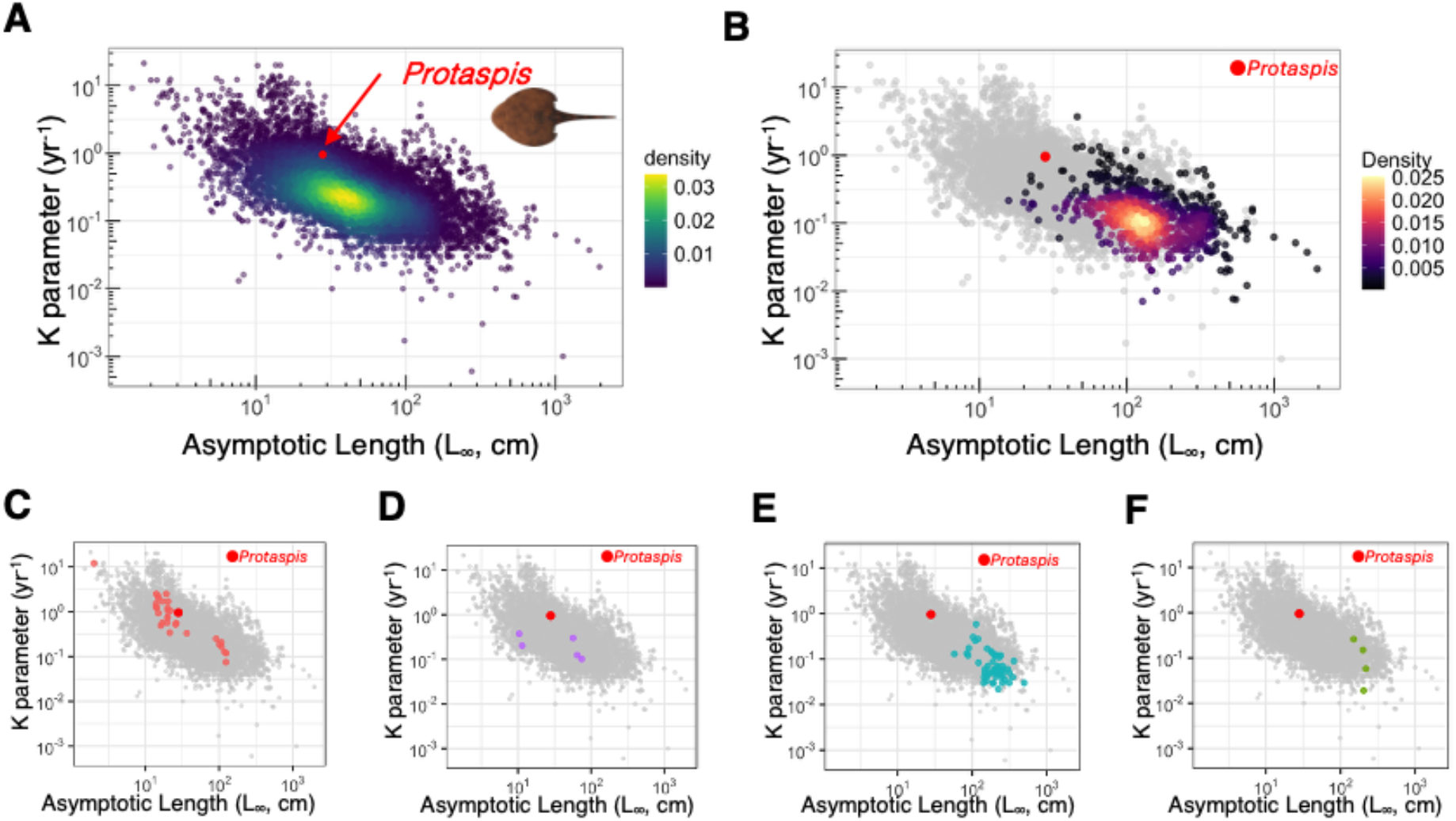
Comparison of asymptotic length and growth parameter *K* between extant fishes, non-Teleostean fishes, armored body fishes and *Protaspis*. Parameters for the corresponding fish were retrieved from rFishBase and compared with Protaspis using a scatter plot. X-axis: Asymptotic Length (L_oo_, cm), Y-axis: Growth parameter K. (A) Extant fish are represented by all data registered in rFishBase, n = 13,374. *Protaspis* is shown as red points and extant fishes are shown as blue to yellow points in the density distribution. (B) Non-Teleostean fishes in rFishBase, n = 766. *Protaspis* is shown as red points and extant fishes are shown as black to yellow points in the density distribution. Other fishes are shown as grey. (C)∼(F) Among extant fish possessing a hard, armored body similar to *Protaspis*, comparisons with *Protaspis* were made based on the elements constituting the armor. (C) Fish with entire bodies covered by bony plates; n = 30, which includes pipefish, cownose ray and seahorses. (D) Fish with scutes; n = 5, including catfish and pinecone fish. Species overlap is ignored; red indicates *Protaspis*, other armored fish are shown in different colors, and all other fish on the rFishBase are shown in gray. (E) Fish with scales made of ganoid; n = 56. Includes Sturgeon and Paddlefish. (F) Fish with scales made of cosmin; n = 6, including coelacanth.

Next, we compared it with Non-teleostean fish, which are among the earliest extant fish (Figure 4B, S8). The center of the Non-teleostean group was L_∞_ = 126 cm and K = 0.12 year^-1^. The group’s plots clustered in the lower right, indicating extremely slow growth and predominantly large bodies. Although phylogenetically closer to *Protaspis*, these extant ‘living fossils’ reach their large size via modest growth rates sustained for a long time. Also, comparisons were made with flat, benthic fish roughly similar morphology and ecology to *Protaspis* (Figure S10). Similar fishes’ density average was nearly identical to that of extant fish overall and the arrangement showed a different center of mass compared to *Protaspis*.

Finally, comparing extant fishes with different types of external armor revealed distinct distributions across the four armor categories (Fig. 4C-F). Fish possessing bony plates included taxa such as pipefish (*Syngnathus* spp.), cownose ray (*Rhinoptera* spp.) and seahorses (*Hippocampus* spp.). These taxa generally occupied the upper left regions of auximetric plots, characterized by low values of L_oo_ and high values of K. Fish with scute type armor were primarily represented by armored catfish (*Rhinelepis* spp.) and pinecone fish (*Monocentris* spp.). Similar to the bony-plate group, these taxa also tended to occupy the upper-left region of the parameter space, characterized by relatively small body size and relatively high K values.

In contrast, fish with cosmin scales, including the coelacanth (*Latimeria chalumnae*.), were distributed in the lower right region of the plot, reflecting larger asymptotic lengths and lower growth coefficients. A similar pattern was observed in fish with ganoid scales, including taxa such as the paddlefish (*Polyodon* spp.), and sturgeon (*Acipenser* spp.). Overall, fishes with cosmin and ganoid scales, predominantly teleosts, tended to cluster in the lower-right portion of the parameter space, whereas fishes with bony plates and scutes, comprising ray-finned fishes, tended to cluster toward the upper-left portion of the distribution, exhibiting a trend similar to that of *Protaspis*. All these comparisons were also performed on all subsets (cwtr, ss, sstr), and it was confirmed that each plot showed relationships indistinguishable from those of *Protaspi*s (main dataset: cw) (Figure S8, S9, S10, S11).

## Discussion

This study combines morphological observations, assessment of plate completeness, and growth modeling using ELEFAN applied to a large fossil dataset to provide the first population-level reconstruction of *Protaspis*’s development and life history strategy. Growth parameters are robustly estimated by ELEFAN because, being non-parametric, it ignores outliers. Here, using ELEFAN suggested rapid early growth with distinct seasonal recruitment, and a short lifespan, implying a fast-paced life history strategy. These characteristics stand in sharp contrast to the long-held assumption that early vertebrates were uniformly slow-growing and long-lived. Quantitative evidence of such life history strategies in Paleozoic vertebrates is novel. Similar strategies are well-documented in extant fish, particularly in environments characterized by seasonal instability and high mortality but were rarely inferred from fossil data. Therefore, these results provide direct evidence that at least some early vertebrates were characterized by rapid growth and high rates of population turnover.

Importantly, the estimated growth parameters showed high consistency across multiple subsets of the dataset, including analyses restricted to *C. transversa* and/or restricted to specimens from the S side of Cottonwood Canyon. The similarity in K-estimates in Table S2 across these subsets attest to the relative robustness of ELEFAN to small differences in taxonomic or geographic sampling. This robustness supports the use of genus-level datasets for reconstructing the growth dynamics of *Protaspis*.

Growth modeling and plate completeness analysis further revealed a distinct developmental separation between body growth and the development of ossified dermal armor in *Protaspis*. Rapid body growth occurred during the juvenile stage, a period when the dermal plates were incomplete and weakly integrated. By reaching a dorsal plate length of approximately 70 mm, plate completeness and fusion increased significantly. This closely corresponds to the size threshold separating 0-year-old and 1-year-old individuals estimated by ELEFAN. This close correspondence strongly suggests that *Protaspis* prioritized rapid body growth during its first year of life, with reinforcement and stabilization of the dermal armor occurring primarily during its second year. This pattern indicates that the presence of dermal armor did not restrict early growth. Rather, the defensive structure appears to have been developmentally delayed, consistent with a trade-off between growth rate and defensive stability—a pattern widely observed in extant fish (Arendt et al., 2001; Barrett, 2010).

The observed increase in plate completeness and adhesion suggests that fusion of the skin plates may have reduced the flexibility of the dermal skeleton. Therefore, juvenile *Protaspis* likely possessed a more adaptable and deformable body surface. Functionally, this state may resemble the flexible scale covering seen in many extant fishes, which facilitates body deformation during swimming and feeding (Vernerey & Barthelat, 2010, 2014). Although heterostracans are not considered aggressive predators (Denison, 1961; Purnell, 2002), increased flexibility during the juvenile stage may have enabled improved swimming efficiency and reduced body compression during suction feeding, even without fully developed jaws (Sanford & Wainwright, 2002). Although direct evidence for feeding behavior is lacking, the inferred flexibility of juvenile facial armor biomechanically supports such a mechanism (Jimenez et al., 2018).

In contrast, adults exhibit a fully movable rigid plate structure accompanied by the development of a characteristic rostrum (Denison, 1967; Janvier, 1996). The mechanical integration of the adult armor likely restricted body deformation and reduced swimming efficiency (Kolmann et al., 2020). This may have facilitated hydrodynamic stabilization via the rostrum (Patel & Riveros, 2013), as proposed for other contemporaneous fish or fish with similar beaks, or a benthic, low-mobility ecological strategy (Ferrón et al., 2020). Although tentative, these functional interpretations are consistent with the observed patterns of ontogeny and growth.

The life history strategy reconstructed for *Protaspis* fundamentally differs from modern slow-growing armored fish, deep-sea species, cartilaginous fish, and non-teleost lineages often treated as ecological analogues for early vertebrates (Mahé et al., 2021; Pikitch et al., 2005; Rigby & Simpfendorfer, 2015). *Protaspis* does not exhibit low growth rates and long lifespans, but rather a combination of high growth coefficients, small asymptotic body sizes, and rapid cohort turnover. This trait combination is well-documented in extant teleosteans and bony-armored fish inhabiting seasonal or unstable environments, where a rapid life history confers a selective advantage (Furness, 2016; Gayanilo Jr et al., 1988; Pecuchet et al., 2017; Reichard & Polačik, 2019; Vrtílek et al., 2018). Therefore, our findings challenge the assumption that extant armored fish, represented by ‘living fossils,’ are appropriate ecological models for early agnathans.

Previous studies have argued that the rapid life history strategy in ray-finned fishes and chondrichthyans evolved as a derived response to environmental instability following ecosystem reorganization in the Late Devonian and Carboniferous periods. Rapid growth, early maturity, and high population turnover rates have been interpreted as evolutionary novelties associated with the radiation of vertebrates after the Devonian (Sallan & Galimberti, 2015). Within this framework, such strategies were not considered characteristic of early vertebrates. Our findings challenge this perspective. The growth rate, cohort structure, and timing of development in *Protaspis* indicate that ecologically fast life histories were already achievable in early jawless vertebrates of the Devonian. Rapid growth and short generation times appear to reflect a widely available ecological strategy that emerges when ecological conditions and mortality rates permit, rather than being a late innovation. Body size is also important for understanding this pattern.

*Protaspis* was significantly smaller than many contemporaneous placoderms, a trait typically associated with high juvenile mortality and rapid life history trade-offs. However, similarly smaller placoderms like *Bothriolepis* have also been found in dense assemblages of juvenile individuals, suggesting high early attrition rates and cohort structuring (Downs et al., 2011; Olive et al., 2016) A similar size-life history relationship is observed in extant cartilaginous fish, where smaller species typically exhibit higher reproductive output and faster life cycles compared to larger relatives (Cortés, 2000).

The contrast between heterostracans, armored fish with bony-covered bodies, and modern, slow-growing armored fish with hard scales, as well as non-teleostean fishes, suggests that conservative growth strategies characterized by low growth rates, large body size, and long lifespans should not be interpreted as ancestral states inherited from early vertebrate evolution (Benton, 2015). Rather, such strategies are likely adaptations to long-term environmental stability (Mahé et al., 2021; Tuljapurkar et al., 2009). The present results indicate that the ‘modern’ rapid-growth life history strategy, characterized by early maturity, short lifespan, and high population turnover, was already established early in vertebrate evolutionary history, long before the origin of jaws or the diversification of crown teleosts. It is worth noting that this strategy appears to be independent of phylogenetic position. When ecological conditions permit, similar life history regimes can evolve repeatedly and independently across vertebrate lineages.

These findings collectively suggest the need to reevaluate the ecology and life history evolution of early vertebrates. Early vertebrates should not be viewed as organisms characterized by persistently slow growth and specialized, robust defense systems across their entire lifespans. Rather, *Protaspis* demonstrates that complex ecological strategies were already present during the early stages of vertebrate evolution, characterized by rapid juvenile growth, delayed stabilization of defense mechanisms, and rapid generational turnover.

This study highlights the extreme importance of fossil population data and quantitative growth reconstruction in elucidating evolutionary strategies that are poorly represented in extant taxa. By combining growth modeling and morphological observations, *Protaspis* emerges as a clear example of a modern bony-armored fish that exhibits rapid growth, a short lifespan, and stage-dependent life history changes. This supports the notion that vertebrate life history evolution was shaped by ecological factors much earlier than previously assumed, rather than being phylogenetically conserved.

## Materials and Methods

### Fossil material and Specimens measurement

The Field Museum in Chicago and the University of Kansas Museum of Natural History in Kansas City, hold a combined collection of over 600 *Protaspis* specimens, which were collected from two sites in the Beartooth Butte Formation of Wyoming: the Beartooth Butte locality and the Cottonwood Canyon locality. In this study, we utilized a total of more than 450 specimens from the Cottonwood Canyon locality, which has a larger specimen collection. This collection yielded the largest number of specimens bearing either the entire dorsal plate or ventral plate, or portions of either (but specimens consisting solely of the orbital plate or rostrum were not used).

This fossil collection contains several species of the genus *Protaspis* from several collection sites within the same locality (Table S1). While this site information is registered in the museum database, specimens registered as “*S side of Cottonwood Canyon*” are listed as “east of Lovell, NE” in Denison (1970), occurrence, making it difficult to determine the exact area represented. According to Fiorillo (2000), Cottonwood Canyon is represented by two distinct sites: the more commonly collected southern wall locality and a rock unit exposed on the extremely difficult-to-access northern wall of the canyon. While the two sites examined in this study were previously thought to be separate areas, based on the information in the original description paper, they may have been collected from the same region.

This study aims to elucidate population structure and seeks to analyze as many specimen groups as possible. Since time-averaged fossil samples exhibit parameters similar to those of populations (Hunt, 2004), they are treated collectively as main dataset. The Cottonwood Canyon Locality contains five *Protaspis* species, and specimens of *Protaspis* sp. which couldn’t be assigned to any species.

The species classified as *Cosmaspis (Protaspis) transversa, Protaspis brevispina, P. mcgrewi, and P. ovatus* were grouped for analysis to increase the sample dataset, due to their similar morphology and similarity in size distribution (Figure S2, S3, S4, S5). Although *P. ovata* tended to exhibit smaller sizes, it was determined that including it in the dataset would not impact the growth analysis, given the inherent robustness of ELEFAN analyses. This conclusion was reached based on the fact that it comprises only five samples; also considering the plate preservation rate, *P. ovata* is considered to exhibit the same juvenile morphology as other species. Additionally, to verify differences in results when multiple sites and species are grouped together, analyses were also conducted using subset: sub dataset cwtr = only *C. transversa* from Cottonwood Canyon locality, sub dataset ss = all specimens from S side of Cottonwood Canyon site, sub dataset sstr= only *C. transversa* from S side of Cottonwood Canyon site (Figure S1).

A total of 455 specimens were examined, ranging from juvenile to adult stages as inferred from body size and the shapes of plates (Figure S2, S3). All specimens were photographed using a Nikon D7500 camera equipped with a Nikon AF-S DX Micro Nikkor 40mm f/2.8G lens and acquired as image data with a scale bar. Specimen photographs were analyzed using ImageJ to measure standard lengths (Schneider et al., 2012). To evaluate potential measurement error, a portion of the sample was measured repeatedly, confirming that the variation was negligible relative to the size class range. The standard length was defined as the longest vertical and horizontal dimensions of either the Dorsal or Ventral plate, measured using ImageJ (Dataset S1). Observation of complete specimens confirmed that the size between Dorsal and Ventral Plates was nearly identical. Therefore, no error between plates was assumed, and measurements proceeded accordingly. Lengths were recorded alongside specimen numbers and other information. To avoid size-related biases in preservation status, measurements were taken and added to the data for specimens where the vertical size could be measured or inferred from the overall shape, even if parts were missing.

### Plate completeness analysis

*Protaspis* consists of distinct plates: Rostrum, Pineal, Orbital, Branchial, Dorsal, Ventral, Dorsal fin, and a Tail (Denison, 1967). Plates are thought to become harder and thicker with age, adhering more closely to other plates. Therefore, examining the preservation rate of each plate allows distinguishing between the active growth stage and the stable, fully grown stage. Since the Dorsal plate has the highest preservation rate and is positioned to attach to other plates, the length of the Dorsal plate was used as the standard to examine the preservation rates of the other plates. The observations were made based on pictures of the specimen using ImageJ. The preservation rate was evaluated on a five-point scale. Based on the total size, the area of each plate was assessed using the following criteria: 5 (100–81%), 4 (80–61%), 3 (60–41%), 2 (40–21%), and 1 (20–1%) (Figure 2, Dataset S1). Plates not identified at all were rated as 0. To prevent observer bias, consistent evaluations were performed by the same observer. The relationship was assessed descriptively.

### ELEFAN analysis

Electronic Length Frequency Analysis (ELEFAN) is a method for detecting cohorts corresponding to distinct spawning seasons from length-frequency (L/F) data, and for estimating the parameters defining growth curves. It is widely used primarily in fishery resource assessment (Pauly & David, 1981). While typically applied to multi-year catch data, it has been demonstrated that useful growth parameters can be estimated from single-year samples when cohort structure is preserved.

This analysis fits the von Bertalanffy Growth Function (Von Bertalanffy, 1938) to length-frequency (L/F) data. The shape of the VBGF is determined by two parameters, asymptotic length (L_∞_), which can be and was here replaced by estimates of maximum length (L_max_) and a growth coefficient (K) which expresses how fast the asymptotic size is approached and which has the dimension 1/time. The VBGF has the form:

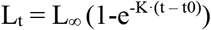

where L_t_ is the length at age t, and t_0,_ which adjust the VBGF to an absolute age scale, is ignored here because it is usually very small, and also because only relative ages are considered here (i.e., t, t+1, t+2, etc.).

ELEFAN superposes thousands of growth curves with different growth parameter combinations onto L/F data restructured using running averages such that their peaks and throughs are well separated and attributed respectively positive and negative ‘points,’ and selects the growth curve as optimal with accumulates most positive points (i.e., best connects the peaks in the L/F data) while avoiding throughs. Its logic is thus based on the assumptions (*i*) that the peaks, when a single sample is examined, represent successive cohorts, and (*ii*) the growth and reproduction/recruitment patterns in the population under study are similar from year to year.

This study conducted length-frequency analysis on 455 specimens of *Protaspis* dorsal and ventral plates collected from the Cottonwood Canyon locality of the Beartooth Butte Formation. Since *Protaspis* dorsal plates and ventral plates are known to be nearly equal in size, with little size variation between species, both were analyzed together without distinction in this study. Furthermore, while *Protaspis* possesses multiple body plates, dorsal and ventral plates exhibit higher preservation rates compared to other plates and are thought to grow continuously during development. Therefore, these plates were used as indicators for growth analysis. Growth parameters were estimated using the TropFishR package in R (Mildenberger et al., 2017) to perform an ELEFAN analysis. First, L/F data were created from *Protaspis* body length measurement data. Since this dataset lacked actual collection time series information, duplicate copies of the same body length data were created and assigned to two pseudo-years (2000 and 2001) to generate L/F data in a format processable by TropFishR. This processing aimed to prepare the data in a computational format suitable for analysis and does not evaluate actual annual variation.

In this study, L_∞_ was fixed at 140 mm for analysis, based on the previously reported maximum length of 137.6 mm for *Protaspis* dorsal/ventral plates.

Next, the growth coefficient K was estimated. Since ELEFAN is an algorithm that tracks the progression of modes on the restructured L/F data, the setting of the length bin size affects the position and sharpness of the peak, thereby influencing the estimated values of K (Wang et al., 2020). Furthermore, fossil data often contains more noise compared to L/F samples for extant fish, potentially obscuring cohort structure. Therefore, this study compared K values estimated for multiple bin sizes rather than relying on a single bin size. Specifically, the body length class interval was varied in 0.1 mm increments within the range of 5.0–20.0 mm. ELEFAN analysis was performed under each condition to estimate K. Moving averages (MA), which are used to restructure the L/F data by smoothing the histogram to facilitate cohort peak detection, must be set to an odd number of bins. In this study, we estimated one cohort width to be approximately 60 mm from the histogram and configured the MA to be input appropriately according to the class interval. The K values obtained for each bin size were plotted against the class interval to identify the region where the estimates were relatively stable. Generally, a bin size that divides length-frequency data into 12–16 classes is considered appropriate. Therefore, this study defined a class interval of 8.7–11.6 mm as the stable region (Figure 3A). The average K value obtained within this range was adopted as the ‘Best K of best K.’ As a result, the representative growth coefficient was estimated as K = 0.95 year^-1^ (standard error = 0.015).

ELEFAN was rerun using the estimated L_oo_ and K, and the resulting growth curves were overlaid on the length-frequency diagram (Figure 3B). Also, VBGF was used to calculate the theoretical length corresponding to each relative age, and the length-frequency histogram was color-coded for each estimated age group (Figure 3C).

Also, to evaluate the impact of dataset bias on estimation results, similar analyses were performed on multiple subsets (Figure S7). Specifically, analyses were conducted using the following as independent datasets: ss = all specimens from all specimens from the S side of the Cottonwood Canyon site, cwtr = only *C. transversa* from the Cottonwood Canyon locality, and sstr = only *C. transversa* from the S side of the Cottonwood Canyon site. The parameters used in each analysis are shown in Table 2.

### Comparing with extant fishes

To compare *Protaspis* growth parameters with those of other fish, we retrieved growth parameters for extant fish species from the rFishBase database and conducted comparative analyses. Analyses were performed using R and the rfishbase (Boettiger et al., 2012) R package. Based on the fact that the dorsal and tail parts are roughly equal in length in other Heterostraci like *Pteraspis* and *Drepanaspis* (Gross, 1963; White, 1935), the maximum body length of *Protaspis* was estimated to be 28 cm and 26 cm. As with the previous growth analysis, four datasets based on different sample groups were used (Table S2) and these were independently overlaid and compared on the distribution maps of extant fish.

First, we compared *Protaspis*’s L_∞_ and K with all extant fish. Using the popgrowth function from the rfishbase package, we obtained estimated values of L_∞_ and K, and ecological information including taxonomic data for all fish species (Dataset S2). Using the log-transformed values of L_∞_ and K obtained, we visualized via auximetric plots, the distribution of growth parameters in extant fish (Figure 4A, S8B, C, D). At this stage, only species possessing both K and L_∞_ were extracted, resulting in a reduced number of species from the original rFishBase dataset. When multiple reported values existed for the same species, the mean values of L_oo_ and K were calculated per species code to obtain representative values at the species level. This averaged dataset was used to analyze the growth distribution across all fish species (Figure S8A). Importantly, we examined all taxonomic groups included in this analysis and excluded fish that rely heavily on air-breathing. The underlying principles of the von Bertalanffy growth function (VBGF) are primarily applicable to fish that obtain oxygen mainly from water through their gills. Taxa that obtain most of their oxygen through air breathing (e.g., *Arapaima gigas*) are known to exhibit growth trajectories that deviate significantly from the typical VBGF shape (Pauly, 1979, 1981). Therefore, to ensure the comparability of growth parameters across species, such taxa were excluded from this analysis. We used a custom dataset based on Graham (1997) and excluded species listed as “Air-breathing” in the Environment field of rFishBase.

Next, we proceed to comparisons with specific taxonomic groups. Using the species names function to obtain the taxonomic class (Class) for each fish species, we classified them into teleost (Teleostei) and non-teleost (other fish), then compared the growth parameter distributions (Figure 4B, S8B, C, D, Dataset S4). Furthermore, using a custom dataset (Dataset S5), parameters for fish with morphologies similar to *Protaspis* and armored bodies were obtained and compared on the plot (Figure S10). Fish with similar morphologies were primarily selected from demersal marine species, chosen for their flat appearance, similar to that of *Protaspis*, based on reference materials such as fish guides. Fish with armor were selected based on having hard bodies or scales like *Protaspis* and were classified into four types: fish with ganoid and cosmin scales, those with scute structures, and those with bodies covered by bony plates. As with other comparisons, parameters were obtained from rFishBase (Dataset S6). The distribution of each group was then displayed on a density plot, with estimated values for *Protaspis*, separated plots for each armored type were created for each of the four structures (Figure 4C, D, E, F, S11).

## Supporting information

This file include Supplementary Text, Supplementary Figure S1-11 and Supplementary Table S1, S2.

## Data availability

The code used to perform the ELEFAN analysis and comparison with extant fishes in this paper, as well as to generate images, is publicly available on GitHub (https://github.com/nanakookabe/Okabe_and_Pauly_2026) and zenodo (https://doi.org/10.5281/zenodo.18162840).

## Acknowledgement

This work was supported by the Sasakawa Scientific Research Grant from The Japan Science Society. We are grateful to many members in the Field Museum and University of Kansas of Natural History Museum for access to specimens. We are grateful to H. Blom (Uppsala University) for valuable suggestions and discussions, and J.N. Wibisana (Okinawa Institute Science and Technology, Graduate School) for coding advice and to Vicky Lam (University of British Columbia) for her input in the ELEFAN analysis. Special thanks go to Franz Anthony (Scientific Illustrator) for his contributions to the reconstructed illustration and 3D modeling.

