## Supplementary material for "Early vertebrates were not slow: rapid life histories in Devonian agnathans": This file include Supplementary Text, Supplementary Figure S1-11 and Supplementary Table S1, S2.

#### Supplementary Information

**Beartooth, first record, environment**

The Beartooth Butte Formation is a stratigraphic unit deposited during the Early Devonian period, located in Wyoming (Bryant, 1932, 1933; Bryant & Ruedemann, 1934; Dorf, 1934). Research into this formation advanced around 1930 following the discovery of vertebrate remains. Sandberg (1961) described this formation as estuarine deposits. However, recent studies using stable isotope analysis suggest it represented a brackish water environment with low salinity and a strong freshwater component. Major fossil localities within this formation include the Beartooth Butte locality and the Cottonwood Canyon locality. Although both belong to the same formation, they are thought to differ in age and depositional environment. Cottonwood Canyon locality is proposed a late Lochkovian to early Pragian age, and Beartooth Butte locality is proposed a mid-late Emsian age (Noetinger et al., 2021). Fiorillo (2000) suggested, based on oxygen and carbon isotope data, that the Beartooth Butte Formation as a whole represented a brackish to freshwater-dominated fluvial and estuarine environment. Within the Formation, the Half Moon Canyon locality shows evidence of potentially stronger freshwater components than other localities, while the Beartooth Butte locality is considered to represent a more marine environment. The Cottonwood Canyon locality is similar to Beartooth Butte, but is considered to have been slightly more brackish.

The Beartooth Butte Formation has reported fossils from numerous biological groups. Vertebrate remains are the most frequently reported. Ostracoderms, primarily Heterostraci and Osteostraci, are well-documented, with the Beartooth Butte Locality showing the greatest diversity, followed by the Cottonwood Canyon locality (Bryant, 1933; Fiorillo, 2000). Many fish of the time had bodies formed from multiple plates, and while most fossils are found with plates detached, Placoderms are more often found with plates still joined. This suggests an environment with moderate flow and water currents capable of detaching plates (Fiorillo, 2000).

Other invertebrate fossils are also present. Lamsdell and Legg (2010) and Lamsdell and Selden (2013) documented the presence of eurypterids and, through classification and phylogenetic analysis, concluded the environment was a relatively warm estuarine or nearshore setting. Furthermore, plant fossils have been discovered. Caruso and Tomescu (2012) reported microconchids forming colonies on the stem sections of terrestrial plant fossils. This indicates that the plants grew partially submerged and that the submerged parts were colonized by microconchids during their lifetime in a freshwater environment, reflecting the conditions at that time. As summarized above, geological and paleontological studies collectively interpret the Beartooth Butte Formation as having formed in a brackish water environment with stronger freshwater components than marine elements. It represents an ancient ecosystem where diverse fish, sea scorpions, and early terrestrial plants coexisted.

**Morphology and ornament rings on the plate of *Protaspis***

*Protaspis* is a genus of heterostracans first described by Bryant (1933). Initially considered part of the genus *Pteraspis,* some species were later reclassified as belonging to the closely related genus *Protaspis*. The genus *Protaspis* is characterized by large dorsal plates and long branchial plates and is believed to have a dorsal shield and ventral shield composed of ten major plates; rostral plate, pineal plate, dorsa plate, ventral plate, left and right orbital plates, parietal plate, and branchial plates. Each plate in the *Protaspis* genus corresponds to these and performs functions such as protecting the head's morphology and forming the area around the respiratory openings. The dermal bone tissue structure of these plates is generally thought to exhibit a four-layer structure: an outer layer of enamel-like-coated dentine projections, a middle layer of acellular parallel bone (the so-called spongy bone layer) containing a vascular network, and a basal layer of lamellar bone, forming an isopedic layer (Keating et al., 2015).

Bryant and Ruedemann (1934) reported that *Protaspis bucheri* and *P. dorfi* had broad dorsal plates and short, broad rostral plates, and slightly broad dorsal plates and symmetrical tail fins, respectively. Subsequently, Denison and Museum (1953) and Denison (1967) described several *Protaspis* species from Water Canyon, Utah. Denison (1970) described the ornamental ridges arranged in plates. These consist of linear or ridge-like dentine projections arranged in plates, with their arrangement differing between species. *P. bucheri* has relatively coarse ornamentation, with ridges running nearly parallel to the growth lines at a density of 4.5–6.5 per mm. The ridges have a smooth, mountain-shaped cross-section, sometimes becoming slightly narrower and granular near the center. *P. dorfi* has extremely fine dorsal ornamentation, with approximately 8–12 fine dentinal ridges per mm running longitudinally across the central region of the scutes. These ridges form sharp, mountain-shaped, unbroken continuous lines. Due to the high ridge density, they are clearly finer than those of *P. bucheri* or *P. macgrewi*. Denison (1970) also reported *P. brevispina, mcgrewi*, and *Cosmaspis (Protaspis) transversa*, used in this study. Specifically: *P. brevispina* has a long, narrow scutum and a long, narrow rostrum, with a finely reticulate ornamentation on the ventral surface of the rostrum. The ornamentation is described as extremely fine, with 8.4 ridges/mm in the central carapace. *P. mcgrewi* has a carapace broader and taller than *P. bucheri* and *P. dorfi*, with a medium-length, broad, rounded rostrum. The carapace surface ridges are somewhat thick and coarse, exhibiting roughly 5.5 ridges/mm in the carapace center. Carapace length (excluding spines) reaches approximately 137 mm. The pattern consists mainly of smooth convex ridges but becomes partially complex with rhombic or small granular elements. The rostrum-ventral region has a well-developed pre-oral area bearing spiny tooth ornamentation. *P. mcgrewi* resembles *P. bucheri* in ornamentation and rostrum shape but is distinguished by the broader scutum and relative length of the rostrum. P. transversa is comparatively large and characterized by banded ornamentation. The ornamentation is convex with smooth apices. The band width is 1.5-2.5 mm, with 4-5 ornaments per millimeter in the lateral direction. This band is arranged concentrically along the growth lines. The body is characterized by a dorsal shield that is wider and taller than that of *P. bucheri*, and a snout that is of moderate length and broad. Based on observation, in *P. transversa*, the rings are formed by elongated oval ornaments arranged side by side (Figure S4). However, on the Dorsal or Ventral Plate, columns composed of vertically elongated ornaments are observed at regular intervals. Examination of well-preserved specimens revealed that these long columns appear at intervals of 10 to 25 columns. While no difference in ornament row thickness was observed near the center, thicker rows appeared toward the outer edges. This was particularly consistent on the Ventral Plate. Furthermore, the shapes of the ornaments forming the rows differed: those in the long rows were elongated, while those immediately before and after were slightly smaller and more closely spaced than the standard ornaments. While this characteristic ornamental row arrangement is most clearly visible in *C. transversa*, which limits our ability to observe numerous specimens, the consistent appearance of thick rows at regular intervals in *C. transversa* suggests that they may function similarly to otoliths, indicating age or growth stage. *P. ovatus* was originally described by Bryant (1933) as *Cyrtaspis ovatus*, but Denison (1970) reclassified it into the genus *Protaspis* due to high morphological similarity. According to Bryant, this species is characterized by a broad branchial plate, a central dorsal plate wider than it is long, and linear ornamentation of beaded projections arranged along the plate's outline, exhibiting a relatively broad, high-bodied shape. *P. ovatus* exhibits a slightly smaller size range compared to other *Protaspis* (Figure S5) but shows similar juvenile morphology as others.

**Ontogeny of *Protaspis***

Although descriptions of the embryology of agnathans, early vertebrates, are extremely limited due to the scarcity and fragility of fossil specimens, reports exist for several species (Cloutier, 2010). Greeniaus and Wilson (2003) detailed the development of the carapace and body scales in the larvae of the Cyathaspididae family of heterostracans, which lived during the same Early Devonian period as *Protaspis*. They reported that the fossils show rapid growth with a unique pattern, where the exoskeleton is sequentially added from the rear to the front and from the body's midline to the sides. Similarly, in Thelodonti, a group of agnathans characterized by a scale-like exoskeleton, squamation initially occurs ventrally and dorsally, progressing along the anterior-posterior axis. Furthermore, observations of scale morphology and growth layers reveal that scales contain multiple generations of odontodes internally, enlarging through the concentric addition of bone and dentine (Chevrinais et al., 2017). This indicates that as the armoring of the head and trunk by large dermal bones progressed, the minute true scales covering the entire body differentiated and developed from dermal odontoderms (Vorob'eva, 2012). In contrast, osteostracans exhibit a pattern opposite to that of heterostracans, where ossification begins laterally on the body and spreads toward the midline (Hawthorn et al., 2008). Furthermore, it has been reported that the shield-like exoskeleton could regenerate and thicken as the animal grew, suggesting that the exoskeleton of extinct agnathans was not merely a rigid shell but biologically active tissue.

*Protaspis* observed in this study, like other Heterostraci and Thelodonti, appeared to grow concentrically from the ossification center (Figure S2, S3, S4). The direction was from posterior to anterior for the dorsal and ventral plates. However, for the rostrum and gill plates, the ossification center was located laterally. While the direction from center to periphery was the same, the anterior-posterior direction differed depending on the plate.

**ELEFAN analysis**

ELEFAN (Electronic Length Frequency Analysis) is a method developed by Pauly and David (1981) to estimate growth parameters from length frequency (L/F) distribution data of fish and other species. It was considered as a growth analysis method for resources where age determination is difficult, such as tropical fish, during the late 1970s and early 1980s. ELEFAN can be used to estimate two of the parameters of the von Bertalanffy Growth Function or VBGF, asymptotic length ($L_\infty$) and K, which expresses how rapidly $L_\infty$ is approached (K is not a 'growth rate' and has the dimension 1/time) by linking the mode of one or several L/F over time with a VBGF (Pauly and David 1981; (Taylor & Mildenberger, 2017).

ELEFAN is widely used by enabling the estimation of growth curves and derived functions (recruitment patterns, mortality, gear selection, etc.) using only readily available L/F readily accessible data compared to other age assessment methods annual rings in hard tissue or tagging-recovery surveys, particularly for tropical fish where such methods are inappropriate or too costly.

While ELEFAN was originally developed as a method for analyzing extant aquatic animals, it has recently been applied to fossil organisms. Thus, Pauly and Holmes (2023) applied ELEFAN to 295 specimens from a fossil assemblages of the trilobite *Triarthrus eatoni* from approximately 450 million years ago, by assuming, as done here for *Protaspis*, that their growth is similar between years. The results showed that trilobite growth followed a von Bertalanffy-type asymptotic growth pattern, rather than the previously assumed linear growth. The estimated asymptotic length $L_\infty$ was 41 mm, and K = 0.29 $year^{-1}$. The estimated lifespan of this trilobite was approximately 10 years, longer than the previously proposed 4 years. The annual mortality rate was also estimated to be around 15–20%, comparable to that of extant crustaceans in the family Peliopidae. This represents the first study to quantitatively evaluate growth parameters for trilobites, and the method is suggested to be applicable to other trilobites and other fossils. In the Supplementary Materials to their paper, Pauly and Holmes (2023) explain why the VBGF, as growth curve, is appropriate for the growth analysis of fossils of small water-breathing organisms.

Similarly, Wu et al. (2024) performed ELEFAN analysis on the predatory radiodont *Amplectobelua symbrachiata*, which lived approximately 520 million years ago during the Cambrian period. These authors analyzed a L/F distribution of 432 fossil specimens using the lengths of one of their claws, ranging from 9.1 to 137.1 mm, and based thereon, confirmed a previous inference (based on the size differences between successive instars) that this animal grew exceptionally fast. This unusually high growth rate is argued to represent an adaptive strategy enabling its success as a top predator in the Cambrian ecosystem, potentially constituting part of the arms race during the Cambrian Explosion.

Also, an ELEFAN analysis has been performed on fossil organisms from the Ediacaran Period (approximately 550 million years ago), known as the earliest multicellular fauna. Ivantsov et al. (2025) analyzed 211 specimens of the Ediacaran organism *Parvancorina minchami* from a single lithofacies in the White Sea region of Russia. The results indicate that the growth of this species fits the VBGF well, with an estimated $L_\infty$ of about 2 cm and K of 1.0 $year^{-1}$. Additionally, the natural mortality coefficient (M) was estimated to be 1.44 $year^{-1}$, with the M/K ratio of 1.4 well within the range observed in extant small invertebrates. From these parameters, the average lifespan of *P. minchami* was estimated to be approximately 4 years. Surprisingly, this Ediacaran organism, whose morphology and phylogenetic position remain unclear, may have shared growth patterns and lifespan characteristics with many extant invertebrates.

These studies suggest that ELEFAN analysis provides a new perspective for understanding Paleozoic organisms ecologically. Indeed, using vast fossil assemblages from a single sub-layer, as in this study of protaspids, represents virtually the only method capable of estimating lifespan from fossils. Furthermore, the ability to infer ecology by comparing growth rates with extant fish represents a powerful approach.

### Figures

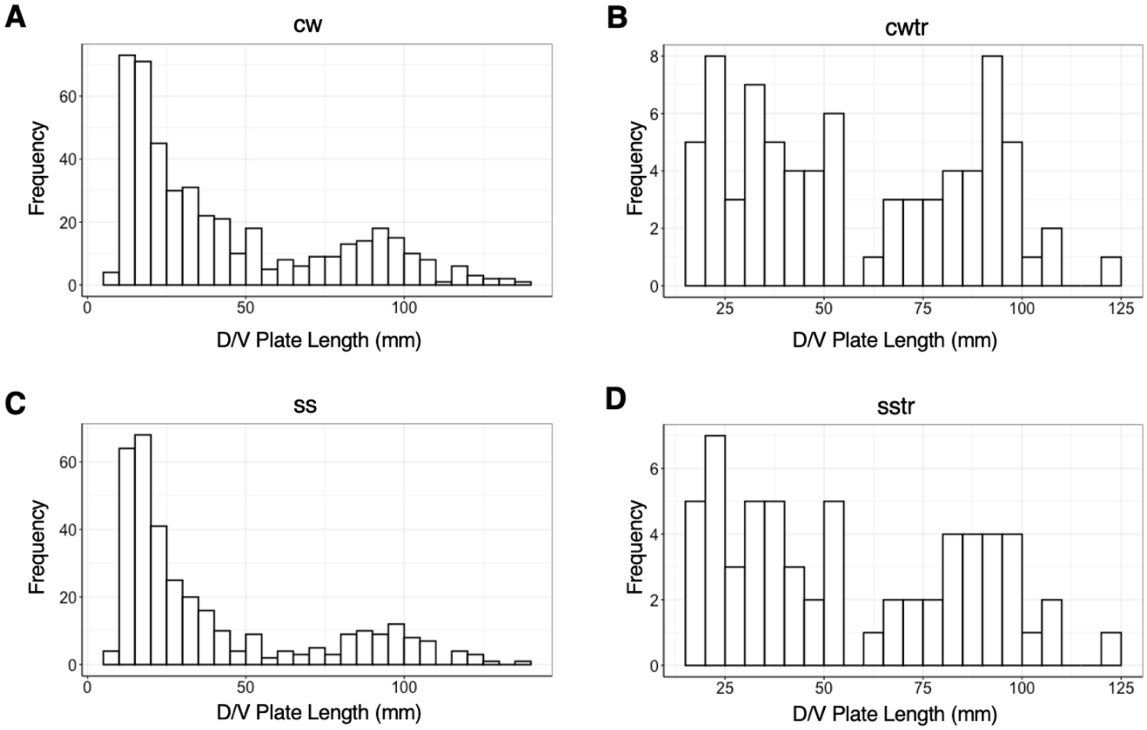


**Fig. S1**. Length/frequency histogram of *Protaspis* for each dataset; Each dataset is represented by a histogram with the X-axis (mm) showing dorsal and ventral plate size bins in 5-millimeter increments and the Y-axis showing frequency. (A) cw: Main dataset containing all specimens from multiple species and sites collected at the Cottonwood Canyon locality; n= 455. (B) cwtr: Histogram of *Cosmaspis* (*Protaspis*) *transversa* specimens collected from two sites within the Cottonwood Canyon locality; n = 77. (C) ss: Histogram including all specimens, across multiple species, cataloged in the museum with Site as S side of Cottonwood Canyon; n = 342. (D) sstr: Histogram of *Cosmaspis* (*Protaspis*) *transversa* specimens cataloged in the museum with Site as S side of Cottonwood Canyon; n = 62.

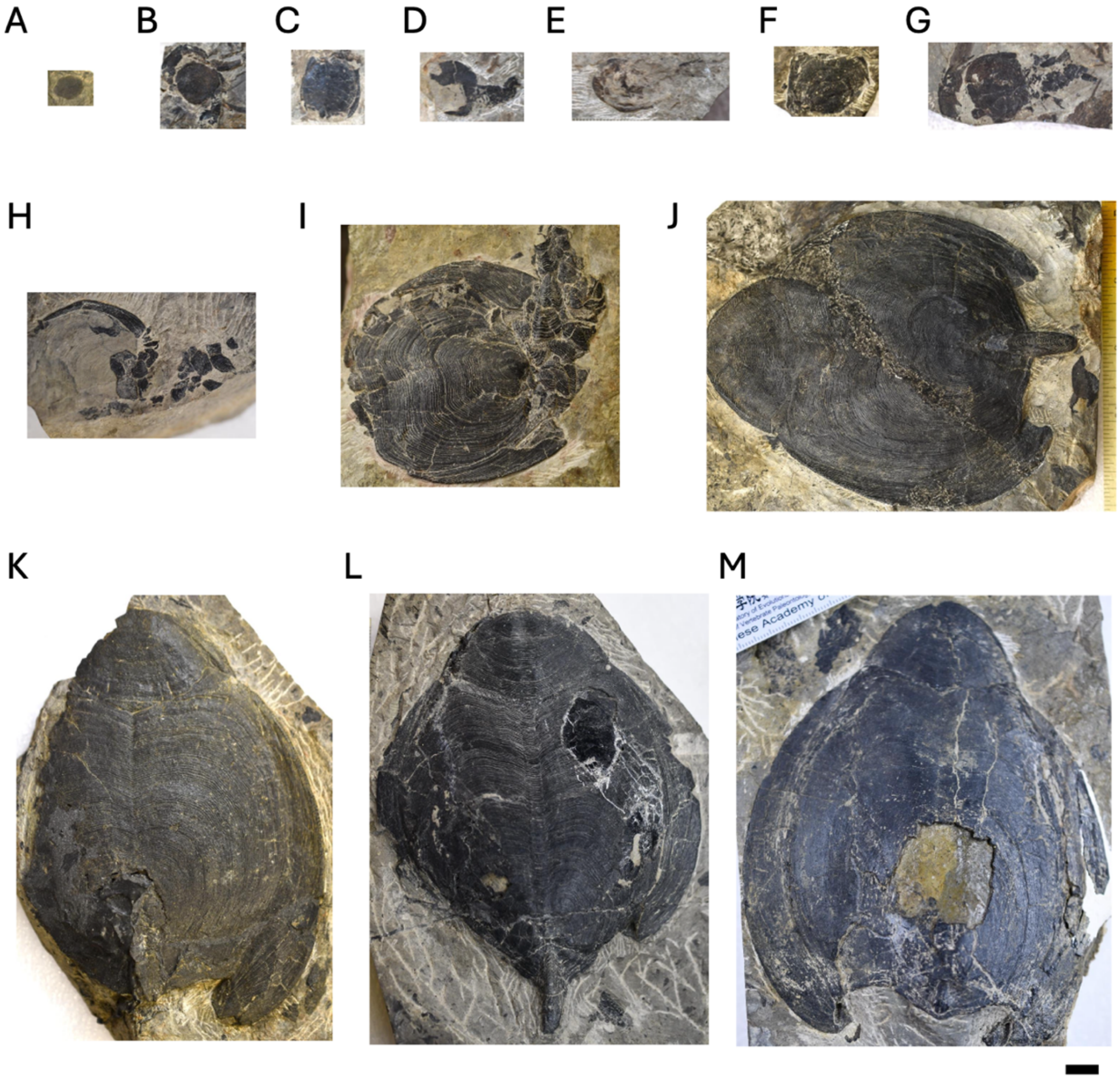


**Fig. S2.**

Fossil specimens of *Protaspis spp.* showing growth series; *Protaspis* spp. specimens arranged by size. All specimens are from the Field Museum collection. The black line at the bottom right is 10 mm. The collection registration names, specimen numbers, and lengths of the Dorsal or Ventral plates for all specimens are shown following. (A) PF5039, 9.6 mm. (B) PF4958, 15.1 mm. (C) PF4491, 15.7 mm. (D) PF4957, 15.8 mm. (E) PF4966, 18.1mm. (F) PF4496, 22.6 mm. (G) PF4494, 23.2 mm. (H)PF4969, 32.2 mm. (I) PF4490, 46.6 mm. (J) PF4334, 53.6 mm. (K) PF4755, 68.6 mm. (L) PF4759, 84.5 mm. (M) PF4340, 97.8 mm.

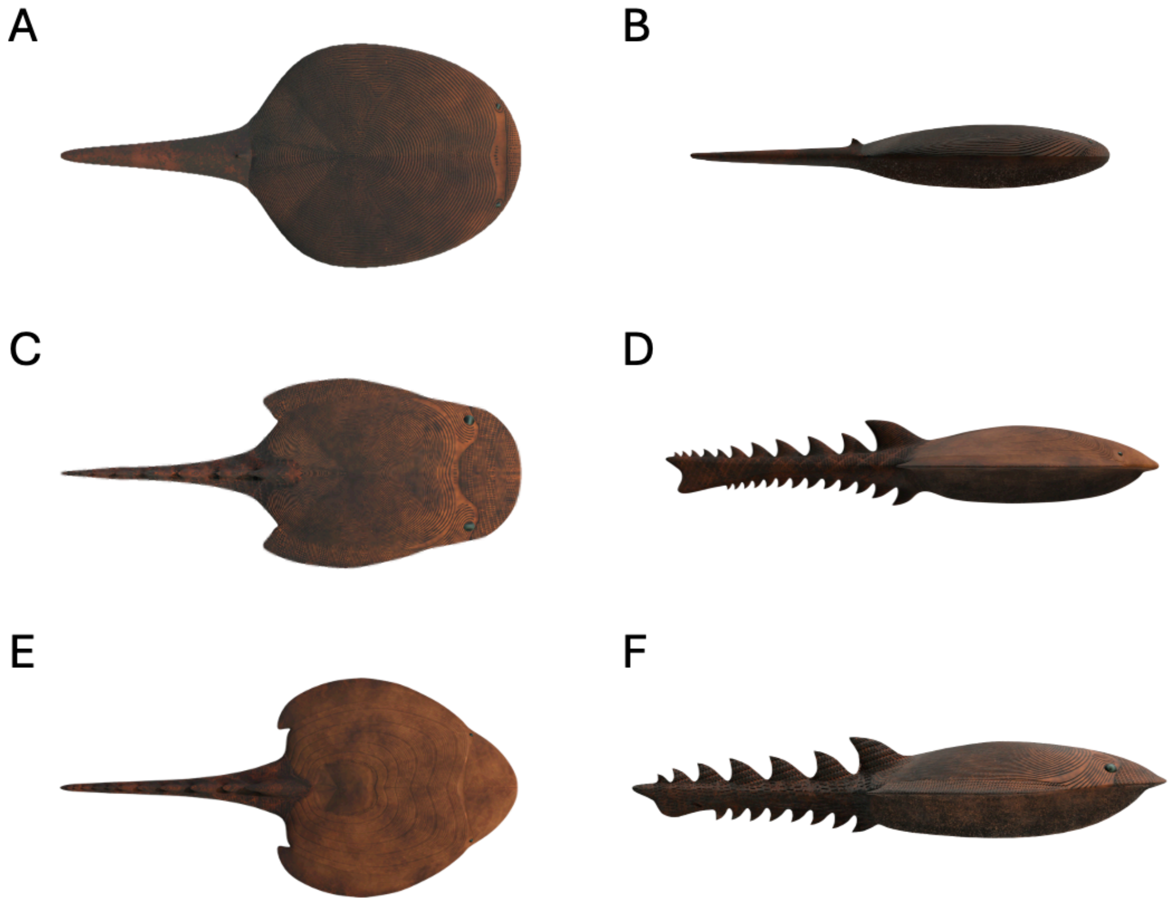


**Fig. S3.** Growth series of *Protaspis;* (A)Juvenile stage 1 viewed from directly above. (B)Juvenile stage 1 viewed from directly beside. (C) Juvenile stage 2 viewed from directly above. (D) Juvenile stage 2 viewed from directly beside. (E) Adult stage viewed from directly above. (F) Adult stage viewed from directly beside.

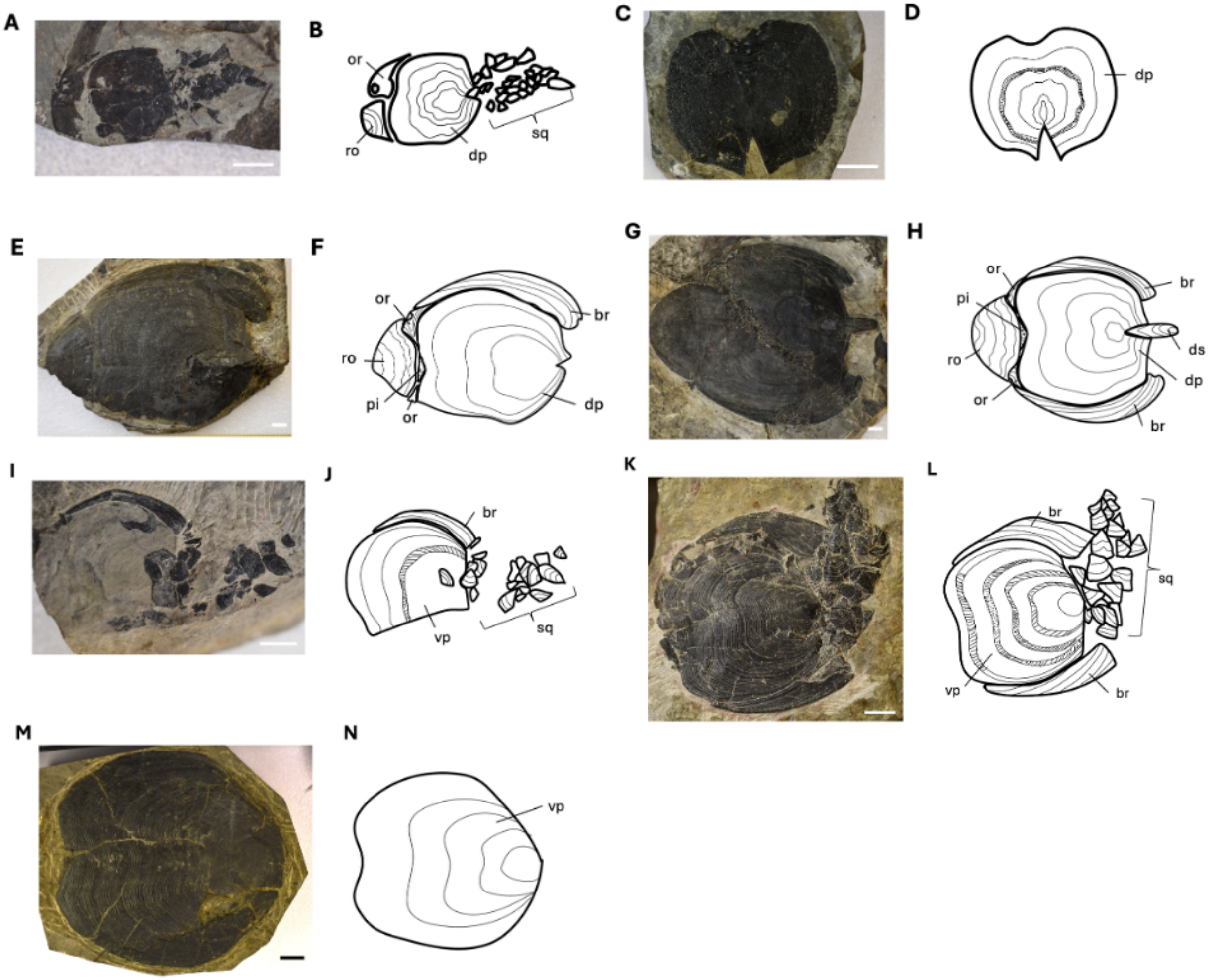


**Fig. S4.** Growth series of *Cosmaspis (Protaspis) transversa*; (A)~(H) Dorsal plate side, and (I)~(N) Ventral plate side. (A, B) PF4494. Dorsal plate length = 23.2 mm. (C, D) PF4984. Dorsal plate length = 41.3 mm. (E, F) PF4755. Dorsal plate length = 68.6 mm. (G, H) PF4334. Dorsal plate length = 89.2 mm. (I, J) PF4969. Ventral plate length = 32.2 mm. (K, L) PF4490. Ventral plate length = 46.6 mm. (M, N) PF4363. Ventral plate length=98.1 mm. or = Orbital plate, ro = Rostral plate, dp = Dorsal plate, sq = Scales, pi = Pineal plate, br = Branchial plate, ds = Dorsal spine, vp = Ventral plate. White bar in specimen photos in (A) - (K) is 1 cm, and black bar in (M) is 2 cm. The line on the dorsal/ventral plate represents a thick ornamental line that appears periodically.

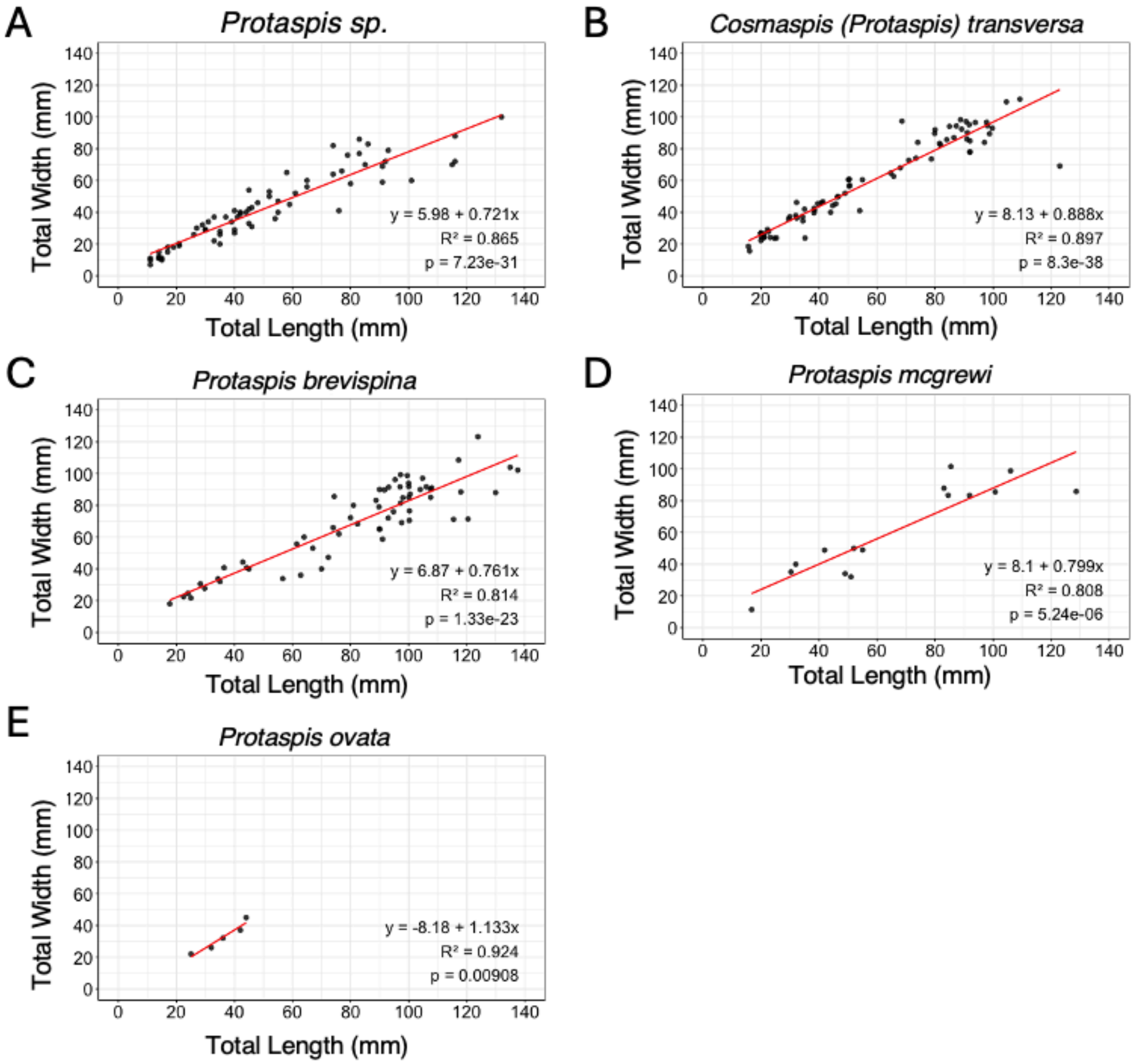


**Fig. S5.** Scatter plot and regression analysis of dorsal/ventral plate size ratios for each *Protaspis* species used in this study; (A) *Protaspis* sp. n = 296. (B) *Protaspis transversa* n = 77. (C) *Protaspis brevispina* n = 62. (D) *Protaspis mcgrewi* n =15. (E) *Protaspis ovatus* n = 5. All plots have the dorsal/ventral plate length (mm) on the X-axis and the dorsal/ventral plate width (mm) on the Y-axis. The red line represents the results of the regression analysis.

**A**

cwtr

| Class (mm) | N= | Dorsal | Rostrum | Orbital | Pineal | Branchial | Spine | Average |
|---|---|---|---|---|---|---|---|---|
| 0-10 | 0 | 0 | 0 | 0 | 0 | 0 | 0 | 0 |
| 10-20 | 5 | 4.80 | 1.00 | 1.00 | 1.00 | 0.40 | 0.00 | 1.37 |
| 20-30 | 9 | 4.67 | 0.56 | 0.78 | 0.22 | 0.56 | 0.22 | 1.17 |
| 30-40 | 5 | 4.00 | 0.00 | 0.00 | 0.00 | 0.00 | 0.00 | 0.67 |
| 40-50 | 4 | 4.25 | 0.00 | 0.00 | 0.00 | 0.00 | 0.00 | 0.71 |
| 50-60 | 5 | 4.60 | 0.00 | 0.00 | 0.00 | 0.00 | 0.00 | 0.77 |
| 60-70 | 3 | 4.33 | 2.67 | 1.33 | 2.33 | 2.00 | 0.67 | 2.22 |
| 70-80 | 4 | 4.25 | 3.25 | 3.25 | 2.75 | 2.75 | 2.25 | 3.08 |
| 80-90 | 8 | 4.25 | 3.63 | 3.63 | 3.88 | 3.25 | 2.25 | 3.48 |
| 90-100 | 10 | 4.40 | 3.00 | 2.90 | 2.60 | 3.40 | 3.30 | 3.27 |
| 100-110 | 2 | 5.00 | 5.00 | 4.50 | 5.00 | 5.00 | 5.00 | 4.92 |
| 110-120 | 0 | 0 | 0 | 0 | 0 | 0 | 0 | 0 |
| 120-130 | 1 | 5.00 | 5.00 | 5.00 | 5.00 | 5.00 | 5.00 | 5.00 |

**B**

ss

| Class (mm) | N= | Dorsal | Rostrum | Orbital | Pineal | Branchial | Spine | Average |
|---|---|---|---|---|---|---|---|---|
| 0-10 | 3 | 4.00 | 0.00 | 0.00 | 0.00 | 0.00 | 0.00 | 0.67 |
| 10-20 | 91 | 4.51 | 0.31 | 0.36 | 0.29 | 0.24 | 0.05 | 0.96 |
| 20-30 | 41 | 4.39 | 0.27 | 0.37 | 0.20 | 0.32 | 0.05 | 0.93 |
| 30-40 | 16 | 4.63 | 0.00 | 0.00 | 0.00 | 0.00 | 0.00 | 0.77 |
| 40-50 | 5 | 4.40 | 0.00 | 0.00 | 0.00 | 0.00 | 0.00 | 0.73 |
| 50-60 | 6 | 4.67 | 0.00 | 0.00 | 0.00 | 0.00 | 0.00 | 0.78 |
| 60-70 | 4 | 5.00 | 1.25 | 1.00 | 1.25 | 1.00 | 1.75 | 1.88 |
| 70-80 | 6 | 4.17 | 2.83 | 2.67 | 2.50 | 2.50 | 1.33 | 2.67 |
| 80-90 | 17 | 3.94 | 3.35 | 3.12 | 3.29 | 2.88 | 2.35 | 3.16 |
| 90-100 | 18 | 4.28 | 4.22 | 4.00 | 4.06 | 3.50 | 2.78 | 3.81 |
| 100-110 | 13 | 4.08 | 4.00 | 3.85 | 4.08 | 3.77 | 2.69 | 3.74 |
| 110-120 | 2 | 5.00 | 2.50 | 2.50 | 2.50 | 2.00 | 2.50 | 2.83 |
| 120-130 | 2 | 5.00 | 2.50 | 2.50 | 2.50 | 2.50 | 2.50 | 2.92 |

**C**

sstr

| Class (mm) | N= | Dorsal | Rostrum | Orbital | Pineal | Branchial | Spine | Average |
|---|---|---|---|---|---|---|---|---|
| 0-10 | 0 | 0 | 0 | 0 | 0 | 0 | 0 | 0 |
| 10-20 | 5 | 4.80 | 1.00 | 1.00 | 1.00 | 0.40 | 0.00 | 1.37 |
| 20-30 | 8 | 4.63 | 0.63 | 0.88 | 0.25 | 0.63 | 0.25 | 1.21 |
| 30-40 | 3 | 4.00 | 0.00 | 0.00 | 0.00 | 0.00 | 0.00 | 0.67 |
| 40-50 | 2 | 4.50 | 0.00 | 0.00 | 0.00 | 0.00 | 0.00 | 0.75 |
| 50-60 | 4 | 4.75 | 0.00 | 0.00 | 0.00 | 0.00 | 0.00 | 0.79 |
| 60-70 | 2 | 5.00 | 2.50 | 2.00 | 2.50 | 2.00 | 1.00 | 2.50 |
| 70-80 | 3 | 4.33 | 3.00 | 2.67 | 2.33 | 2.67 | 1.67 | 2.78 |
| 80-90 | 8 | 4.25 | 3.63 | 3.63 | 3.88 | 3.25 | 2.25 | 3.48 |
| 90-100 | 7 | 4.43 | 4.29 | 4.14 | 3.71 | 3.86 | 3.57 | 4.00 |
| 100-110 | 2 | 5.00 | 5.00 | 4.50 | 5.00 | 5.00 | 5.00 | 4.92 |
| 110-120 | 0 | 0 | 0 | 0 | 0 | 0 | 0 | 0 |
| 120-130 | 1 | 5.00 | 5.00 | 5.00 | 5.00 | 5.00 | 5.00 | 5.00 |

**Fig. S6.** Plate completeness by sub dataset**;** (A) Sub dataset: cwtr. The average plate completeness value increases when the dorsal plate length (class) reaches approximately 60–70 mm. Since there were no samples in the 110–120 mm class, all values are 0. (B) Sub dataset: ss. The average plate completeness value increases around the time the dorsal plate length (class) reaches 60–80 mm. (C) Sub dataset: sstr. The average plate completeness value increases gradually around the time the dorsal plate length (class) reaches 60–70 mm. Since there were no corresponding samples in the 110–120 mm class, all values are 0. Class refers to dorsal plate length (mm). N= indicates the number of corresponding samples. Dorsal, Rostrum, Orbital, Pineal, Branchial, and Spine represent the completeness of each respective plate. Refer to Fig. 2A for the evaluation method. Average denotes the mean value of the completeness scores for these plates.

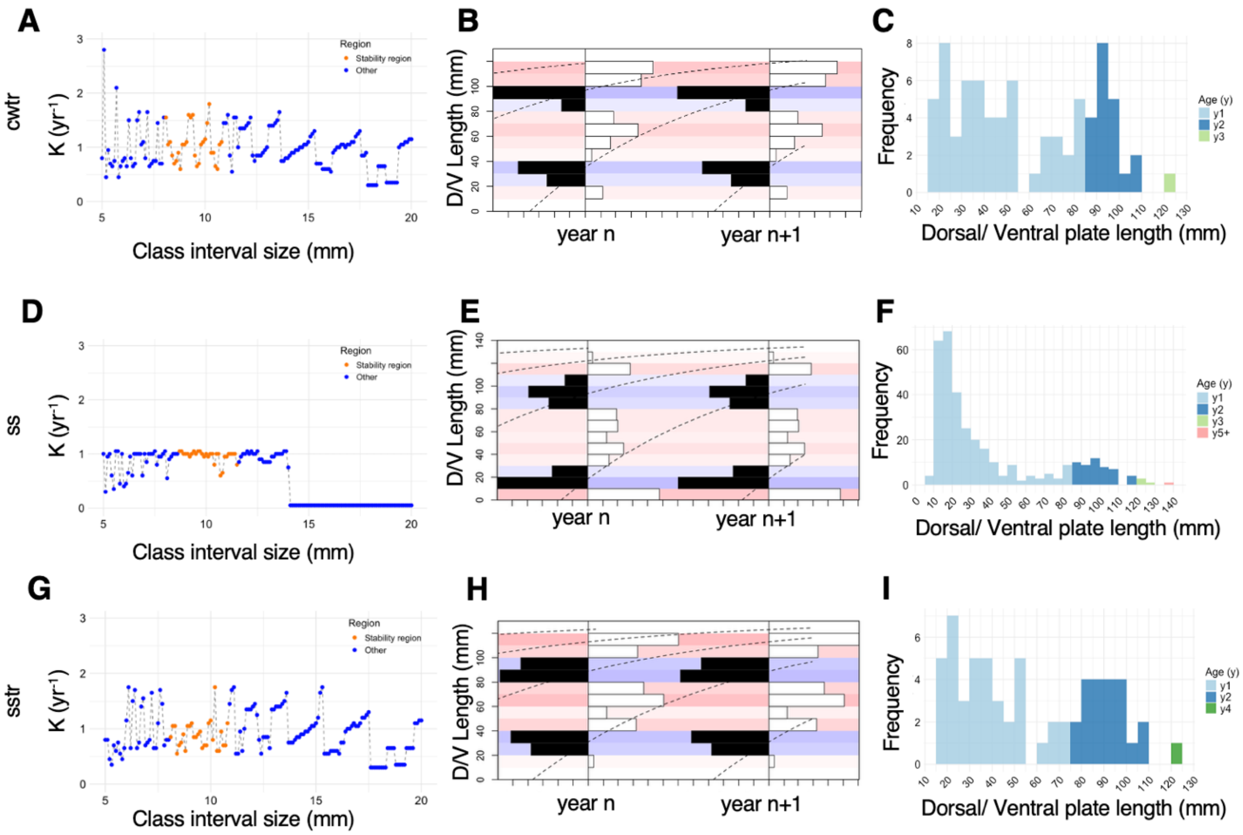


**Fig. S7.** ELEFAN result of each sub dataset; All format and X, Y axis are same as Figure 3. (A) Best K searching for cwtr dataset. Average K (orange dot) is 1.05 year$^{-1}$. (B) ELEFAN plot for cwtr. (C) Age-specific size data for *Protaspis* population structure for cwtr. (D) Best K searching for ss dataset. Average K (orange dot) is 0.96 year$^{-1}$. (E) ELEFAN plot for ss. (F) Age-specific size data for *Protaspis* population structure for ss. (G) Best K searching for cwtr dataset. Average K (orange dot) is 0.88 year$^{-1}$. (H) ELEFAN plot for cwtr. (I) Age-specific size data for *Protaspis* population structure form cwtr.

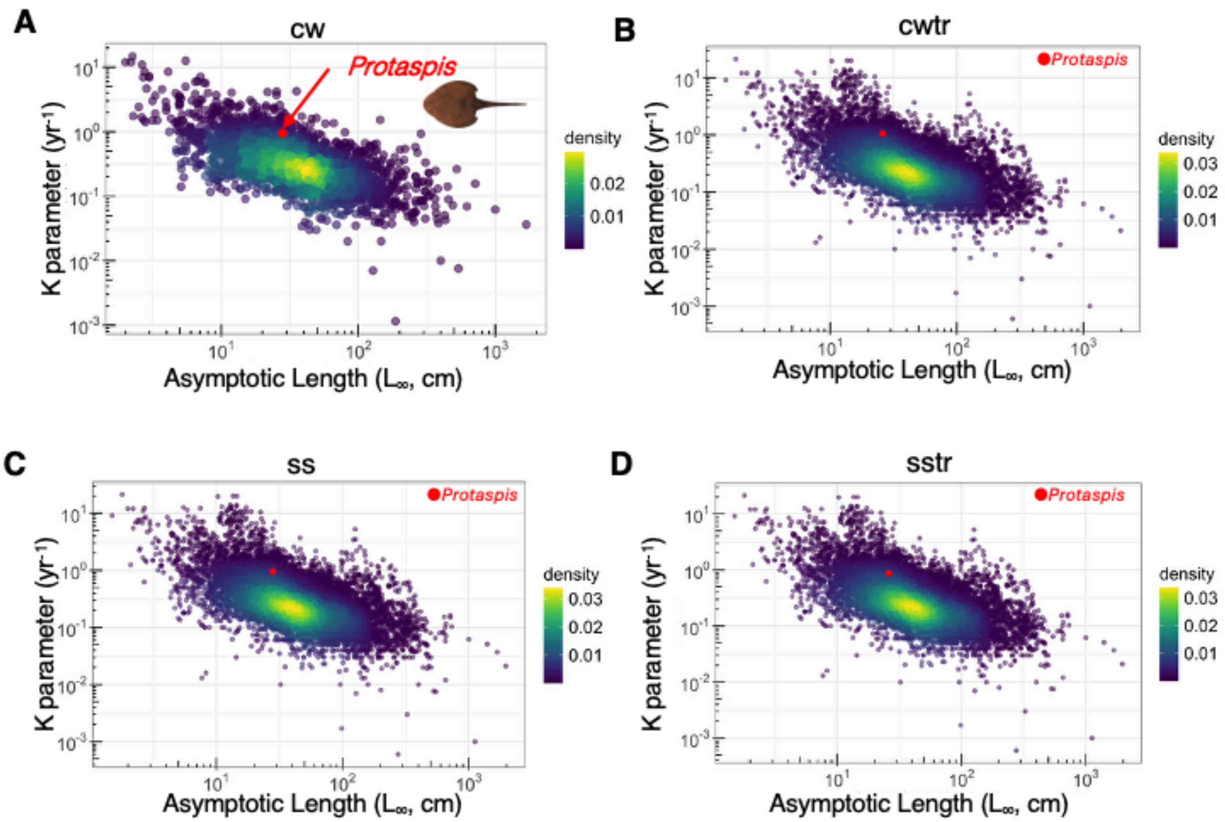


**Fig. S8.** Auximetric comparison of asymptotic length and growth parameter K between extant fish and *Protaspis*. (A): Comparison with species-specific averages excluding duplicate species data; n = 2686. The parameter for this dataset (cw) are $L_\infty$ = 28 cm and K = 0.95 year$^{-1}$. (B) to (D): Comparison with all extant fish species registered in FishBase, including duplicates; n=13374. (B): Comparison with subdataset cwtr: $L_\infty$ = 26 cm, K = 1.05 year$^{-1}$. (C): Comparison with subdataset ss: $L_\infty$ = 28 cm, K = 0.96 year$^{-1}$. (D) Comparison with subdataset sstr: $L_\infty$ = 26 cm, K = 0.88 year$^{-1}$. X-axis: asymptotic length (cm), Y-axis: growth parameter K (year$^{-1}$). *Protaspis* is shown as red points and extant fishes are shown as blue to yellow points in the density distribution.

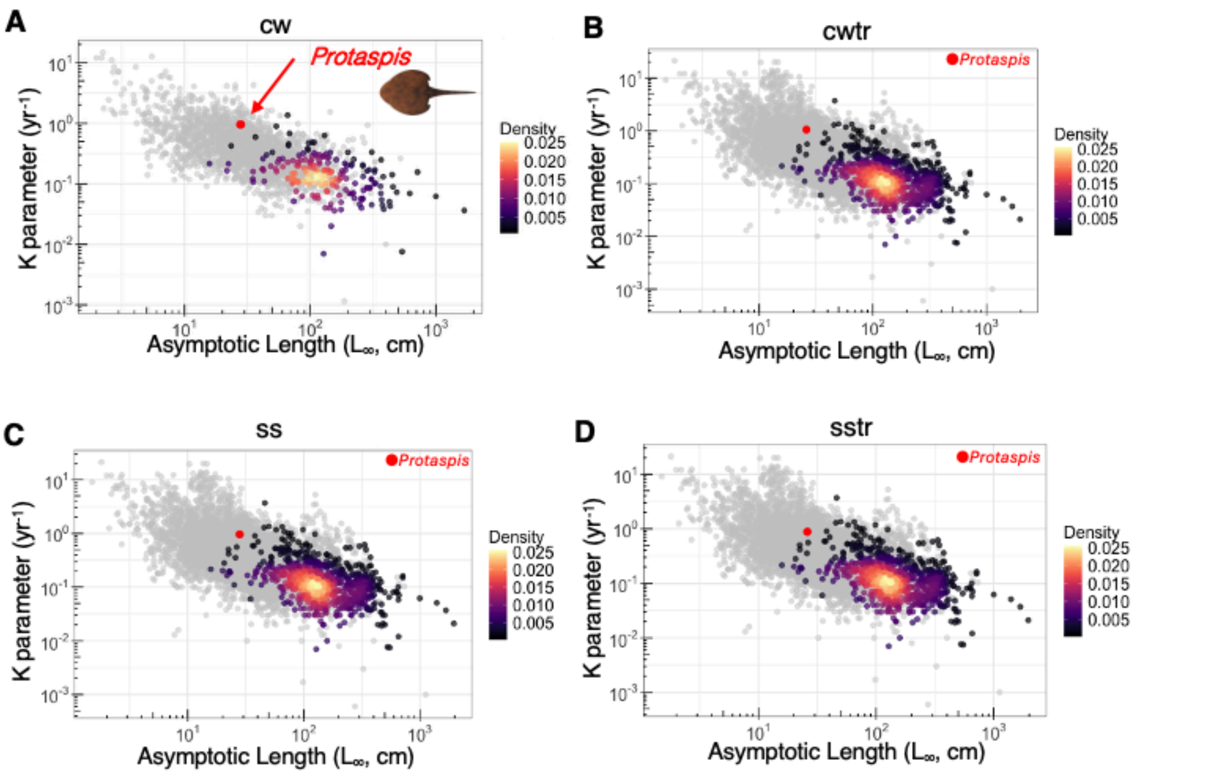


**Fig. S9.** Auximetric comparison of *Protaspis*' asymptotic length and K parameter with non-Teleostean fishes from FishBase. (A): Comparison with non-Teleosteans' species-specific averages excluding duplicate species data; n = 196. The growth parameters for the cw dataset are $L_\infty$ = 28 cm and K = 0.95 year$^{-1}$. (B) to (D): Comparison with all extant fish species registered in FishBase, including duplicates; n=766. (B): Comparing with sub dataset cwtr: $L_\infty$ = 26 cm and K = 1.05 year$^{-1}$. (C) Comparison with sub dataset ss: $L_\infty$ = 28 cm and K = 0.96 year$^{-1}$. (D): Comparison with sub-dataset sstr: $L_\infty$ = 26 cm and K = 0.88 year$^{-1}$. X-axis: Asymptotic length (cm); Y-axis: growth parameter K (in year$^{-1}$). *Protaspis* is shown as red points and extant fishes are shown as black to yellow points in the density distribution.

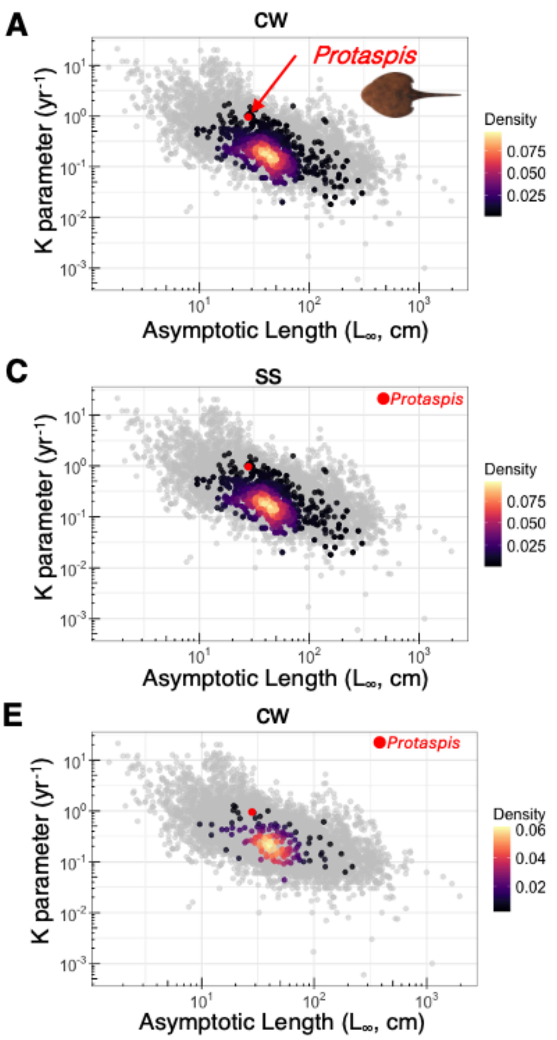


**Fig. S10.** Auximetric comparison of *Protaspis*' asymptotic length and K parameter with fishes similar in morphology to *Protaspis* from FishBase. (A): Comparison with fishes similar in morphology to *Protaspis*'s species; specific averages exclude duplicate species data; n = 132. Parameter of dataset of cw is $L_\infty$ = 28 cm and K = 0.95 year$^{-1}$. (B) to (D): Comparison with all extant fish species registered in FishBase, including duplicates; n = 1101. (B): Comparing with sub dataset cwtr: $L_\infty$ = 26 cm and K = 1.05 year$^{-1}$. (C): Comparison with sub-dataset ss: $L_\infty$ = 28 cm and K = 0.96 year$^{-1}$. (D) Comparison with sub-dataset sstr: $L\infty$ = 26 cm and K = 0.88 year$^{-1}$. X-axis: asymptotic length (cm); Y-axis: growth parameter K (year$^{-1}$). *Protaspis* is shown as red points and extant fishes are shown as black to yellow points in the density distribution.

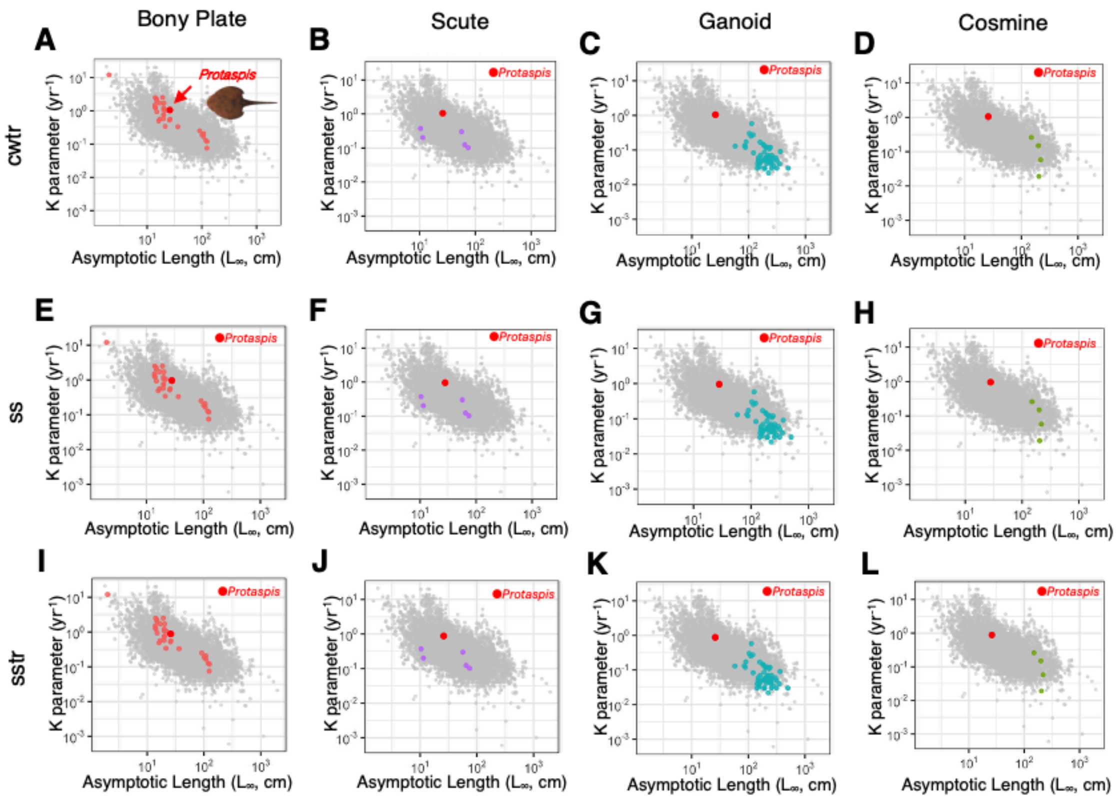


**Fig. S11.** Auximetric comparison of parameters for different types of extant fish with hard armored bodies similar to *Protaspis* and parameters for the *Protaspis* subset; Extant fish with armored bodies similar to *Protaspis* are shown in different colored dots representing different types of armors. (A, E, I) Bony plate: Includes pipefish, cownose ray and seahorse; n = 30. (B, F, J) Scute: Includes catfish and pinecone fish; n = 5. (C, G, K) Ganoid: Includes sturgeon and paddlefish; n = 56. (D, H, L) Cosmine: Includes coelacanth; n = 4. The *Protaspis* parameters obtained for each subset are as follows. (A, B, C, D) cwtr: $L_\infty$ = 26 cm, K = 1.05 $year^{-1}$. (E,F,G,H) ss: $L_\infty$ = 28 cm, K = 0.96 $year^{-1}$. (I, J, K, L) sstr: $L_\infty$ = 26 cm, K = 0.88 $year^{-1}$. All histograms show X-axis: Asymptotic length (cm), Y-axis: Growth parameter K ($year^{-1}$).

### Tables

**Table S1.** List of *Protaspis* specimens collected in the Field Museum of Natural History and the KU Natural History Museum. The Beartooth Butte Formation includes two localities: the Beartooth Butte locality contains 44 specimens, and the Cottonwood Canyon locality contains 455 specimens. Each locality also includes multiple collection sites, the locations of which are based on information from the museum's database. '*Protaspis* sp.' includes mainly small individuals where identification to the species level is not possible, and thus 'spp.' would be more appropriate. However, the museum registration name 'sp.' is used here. Only specimens collected by the Field Museum of Natural History and KU Natural History Museum and accompanied by whole or partial dorsal or ventral plates, were used, i.e., specimens consisting solely of orbital or rostrum plates were not used. This study uses only specimens collected primarily from the Cottonwood Canyon locality. All specimens from Cottonwood Canyon form the cw and cwtr dataset, while specimens collected from the S side of the Cottonwood Canyon site form the ss and sstr dataset.

| Location | Site | Species name | Number of specimens |
|---|---|---|---|
| Beartooth Butte | Beartooth Butte | *Protaspis (include sp.)* | 16 |
| | | *Protaspis bucheri* | 8 |
| | | *Protaspis ovatus* | 4 |
| | | *Protaspis dorfi* | 3 |
| | | *Protaspis sculptus* | 4 |
| | | Total | 35 |
| | Park county, Northern part of, Outcrop above Beartooth Butte Lake, Less than 5 miles of Montana boundary | *Protaspis ovatus* | 1 |
| | | *Protaspis sculptus* | 6 |
| | | Total | 7 |
| | Upper Bighorn Bear Tooth Buttes | *Protaspis bucheri* | 2 |
| | | Total | 2 |
| Cottonwood Canyon | Cottonwood Canyon, E of Lovell | *Protaspis (include sp.)* | 69 |
| | | *Cosmaspis (Protaspis) transversa* | 15 |
| | | *Protaspis brevispina* | 19 |
| | | *Protaspis mcgrewi* | 5 |
| | | *Protaspis ovatus* | 5 |
| | | Total | 113 |
| | S side of Cottonwood Canyon | *Protaspis (include sp.)* | 227 |
| | | *Cosmaspis (Protaspis) transversa* | 62 |
| | | *Protaspis brevispina* | 43 |
| | | *Protaspis mcgrewi* | 10 |
| | | Total | 342 |

**Table S2.** The *Protaspis* datasets used in this study. The main dataset cw used in this study contains specimens of multiple species collected from the Cottonwood Canyon locality. Subdataset1 cwtr includes all *Cosmaspis* (*Protaspis*) *transversa* specimens collected from the Cottonwood Canyon locality. Subdataset2 ss is a dataset containing multiple species collected from the S side of the Cottonwood Canyon site within the Cottonwood Canyon locality. Subdataset3 sstr is a dataset containing *Cosmaspis* (*Protaspis*) *transversa* collected from the S side of the Cottonwood Canyon site within the Cottonwood Canyon locality. Parameters used in ELEFAN: $L_\infty$ (mm) and K. Parameters obtained from ELEFAN: K and its Standard error. Additionally, sizes estimated from the parameters for each age: y2 size = size at age 1, y3 size = size at age 2, y4 size = size at age 3.

| **Dataset** | **Main Dataset** | **Subdataset 1** | **Subdataset 2** | **Subdataset 3** |
|---|---|---|---|---|
| **Abbrevlation** | cw | cwtr | ss | sstr |
| **N=** | 455 | 77 | 342 | 62 |
| **Location** | | | | |
| COTTONWOOD CANYON, E OF LOVELL | × | × | | |
| S side of Cottonwood Canyon | × | × | × | × |
| **Site** | | | | |
| *Protaspis sp.* | × | | × | |
| *Cosmaspis (Protaspis) transversa* | × | × | × | × |
| *Protaspis brevispina* | × | | × | |
| *Protaspis mcgrewi* | × | | × | |
| *Protaspis ovatus* | × | | × | |
| **$L_\infty$** | 140 | 130 | 140 | 130 |
| **K** | 0.95 | 1.05 | 0.96 | 0.88 |
| **Standard error** | 0.015 | 0.064 | 0.021 | 0.047 |
| **y2 size** | 86 | 84.5 | 86.4 | 76.2 |
| **y3 size** | 119.1 | 114.1 | 119.5 | 107.7 |
| **y4 size** | 131.9 | 124.4 | 132.1 | 120.8 |

#### List of Dataset (Separated file)

**Dataset S1 (separate file).** *Protaspis*_dataset. List of *Protaspis*'s from Cottonwood Canyon locality, size and plate completeness score.

**Dataset S2 (separate file).** Growth_full. Output file of $K/L_\infty$ score for every extant fishes in rfishbase.

**Dataset S3 (separate file).** Air_breathing. Input file of air-breathing fishes**.**

**Dataset S4 (separate file).** Non_teleostei_output. Output file of $K/L_\infty$ score for non-Teleostei.

**Dataset S5 (separate file).** FishbaseComparing_armored_morphology_input. Input file of similar morphology fishes list.

**Dataset S6 (separate file).** Armored_data_output. Output list of $K/L_\infty$ score and armored type of selected armored fishes.
